# Network-based identification and pharmacological targeting of host cell master regulators induced by SARS-CoV-2 infection

**DOI:** 10.1101/2021.07.03.451001

**Authors:** Pasquale Laise, Megan L. Stanifer, Gideon Bosker, Xiaoyun Sun, Sergio Triana, Patricio Doldan, Federico La Manna, Marta De Menna, Ronald B. Realubit, Sergey Pampou, Charles Karan, Theodore Alexandrov, Marianna Kruithof-de Julio, Andrea Califano, Steeve Boulant, Mariano J. Alvarez

**Affiliations:** DarwinHealth Inc, New York, NY, USA; Department of Systems Biology, Columbia University Irving Medical Center, New York, NY, USA; Department of Infectious Diseases, Molecular Virology, Heidelberg University Hospital, Germany; Structural and Computational Biology Unit, European Molecular Biology Laboratory, Heidelberg, Germany; Collaboration for joint PhD degree between EMBL and Heidelberg University, Faculty of Biosciences, Germany; Department of Infectious Diseases, Virology, Heidelberg University Hospital, Germany; Research Group “Cellular Polarity and Viral Infection”, German Cancer Research Center (DKFZ), Heidelberg, Germany; Department for BioMedical Research, Urology Research Laboratory, University of Bern, Bern, Switzerland; Translational Organoid Models, Department for BioMedical Research, University of Bern, Bern, Switzerland; Bern Center for Precision Medicine, University of Bern and Inselspital, Bern, Switzerland; Department of Urology, Inselspital, Bern University Hospital, Bern, Switzerland; Skaggs School of Pharmacy and Pharmaceutical Sciences, University of California San Diego, La Jolla, CA, USA; Molecular Medicine Partnership Unit (MMPU), European Molecular Biology Laboratory, Heidelberg, Germany; Herbert Irving Comprehensive Cancer Center, Columbia University Irving Medical Center, New York, NY, USA; Department of Medicine, Columbia University Irving Medical Center, New York, NY, USA; Department of Biochemistry & Molecular Biophysics, Columbia University Irving Medical Center, New York, NY, USA; Department of Biomedical Informatics, Columbia University Irving Medical Center, New York, NY, USA

**Keywords:** Coronavirus, Regulatory networks, Master regulator, Antiviral drugs

## Abstract

Precise characterization and targeting of host cell transcriptional machinery hijacked by SARS-CoV-2 remains challenging. To identify therapeutically targetable mechanisms that are critical for SARS-CoV-2 infection, here we elucidated the Master Regulator (MR) proteins representing mechanistic determinants of the gene expression signature induced by SARS-CoV-2. The analysis revealed coordinated inactivation of *MR-proteins* linked to regulatory programs potentiating efficiency of viral replication (*detrimental host MR-signature*) and activation of *MR-proteins* governing innate immune response programs (*beneficial MR-signature*). To identify MR-inverting compounds capable of rescuing activity of inactivated host MR-proteins, without adversely affecting the beneficial MR-signature, we developed the ViroTreat algorithm. Overall, >80% of drugs predicted to be effective by this methodology induced significant reduction of SARS-CoV-2 infection, without affecting cell viability. ViroTreat is fully generalizable and can be extended to identify drugs targeting the host cell-based MR signatures induced by virtually any pathogen.

## Introduction

Although the global death toll from COVID-19 is now approaching 3.8M, with almost 152M documented cases worldwide(1), only 2.85 billion vaccine doses have been administered as of June 26^th^, 2021. Problematically, their distribution has been characterized by striking differences based on geography, income, education, and other factors(2, 3). These data highlight the need for a global, multi-pronged approach to complement vaccine-mediated prevention with novel pharmacologic treatments for infected individuals, especially among those at higher risk for poor outcomes. This necessity is strongly supported by the continued emergence of SARS-CoV-2 variants with increased transmission rates and increased risk of resistance to current vaccines. Indeed, the epidemiological community acknowledges that achieving target herd immunity thresholds, estimated to be about 60% to 70% of the population, is unlikely in the foreseeable future.

While the most desirable therapies are those that could be prescribed at the first sign of infection—especially using oral drugs in an outpatient setting—the development and identification of effective pharmaceutical agents to moderate the clinical effects of COVID-19 during early stages of infection remain elusive. In addition to SARS-CoV-2, emergence of new pathogens with pandemic potential is constantly increasing due to population growth and mobility. As a result, availability of novel, rapidly deployable, and pathogen-agnostic methodologies for rapid identification of pharmacological agents that mitigate disease morbidity and mortality associated with viral infection would represent a critical advance in our response to both the current as well as future pandemics.

Several approaches have been developed to identify specific host pathways and proteins whose individual interaction with viral proteins is either required to mediate SARS-CoV-2 infection or represents a key modulator of virulence(4–9). In contrast, there has been relatively little focus on experimental elucidation of host cell transcriptional control mechanisms and programs promoting a pro-viral cellular environment, including identification of Master Regulator (MR) proteins representing viral infection-mediated determinants of the transcriptional regulatory programs hijacked by viruses to improve their replication efficiency and overall infectivity. To address this challenge systematically, we leveraged computational methodologies originally developed in the field of oncology to identify MR proteins controlling the transcriptional state of cancer cells(10) and to prioritize MR-inverting drugs to decommission the regulatory programs required by cancer cells to maintain their aberrant cell state(11). Here we argue that extension and translation of these methodologies to viral infection can identify host cell MR proteins representing key mechanistic dependencies of virtually any viral pathogen, as well as drugs that achieve their therapeutic potential by modulating the activity of these MRs.

Based on their definition, MRs can be accurately and systematically identified by assessing the enrichment of their transcriptional targets in differentially expressed genes, using the Virtual Inference of Protein activity by Enriched Regulon (VIPER) analysis(12). While many approaches can be used to identify the tissue-specific targets of a protein, the Algorithm for the Accurate Reconstruction of Cellular Networks (ARACNe)(13) is one of the few that has been extensively experimentally tested, with validation rates exceeding 70%(13–15). We have shown that VIPER can accurately measure the activity of >70% of regulatory proteins, including in single cells, where we have shown that metaVIPER(16)—a VIPER extension specifically designed for single-cell analyses—can virtually eliminate the gene dropout issue due to low single cell profiling depth(17, 18) and outperform antibody based measurements(17). Both algorithms have been highly effective in elucidating protein-based mechanisms that were virtually undetectable by gene expression-based methods alone(10, 17, 19, 20) (see methods for additional details). Once MR proteins are identified by VIPER analysis, we have shown that the Clinical Laboratory Improvement Amendments (CLIA)-certified OncoTreat algorithm(11) can accurately and efficiently identify small molecule inhibitors that can invert their activity thus collapsing the regulatory programs they control. The latter leverages large-scale gene expression profiles of MR-matched cell lines perturbed with a comprehensive repertoire of clinically-relevant drugs, including Food and Drug Administration (FDA)-approved and late-stage experimental ones and has led to several clinical trials (NCT02066532, NCT02632071, and NCT03211988, among others).

It is therefore reasonable to ask whether these methodologies can be successfully generalized to predict therapeutic agents for non-cancer-related diseases, including those caused by infectious pathogens. Given the urgency mandated by the current COVID-19 pandemic, we decided to test the applicability of this approach to SARS-CoV-2. Specifically, we asked whether these methodologies could be used to identify MR proteins in host cells representing the mechanistic determinants of the transcriptional programs hijacked by the virus to support efficient replication and dissemination, as well as the drugs capable of inverting their activity. Should this prove successful, the methodology could be trivially generalized to other pathogens, conditional only on the availability of appropriate infection gene expression signatures.

VIPER-inferred MRs from multiple SARS-CoV-2 infection models consistently showed that innate immune response programs were primarily governed by host MRs that were *significantly activated* in response to SARS-CoV-2 infection. In sharp contrast, the transcriptional programs required by the virus to achieve optimal replication and infectivity during the hijack phase were controlled by a different repertoire of host MRs that were *significantly inactivated* following infection. We designed the ViroTreat algorithm to identify compounds targeting and counteracting the subset of host cell regulatory mechanisms that are hijacked by the virus to create a pro-infective state, thereby inducing host cell-dependent “viral regulatory contraception”. When used to prioritize a set of 154 FDA-approved drugs primarily indicated in oncology, ViroTreat predictions were highly effective, resulting in >80% success rate in reducing SARS-CoV-2 infectivity in colon epithelial cells, without affecting cell viability.

Based on these findings, we conclude that ViroTreat is not only potentially valuable for identifying therapies in the setting of COVID-19, but this approach can easily be generalized for virtually any pathogen in order to target host cell regulatory mechanisms that facilitate viral hijacking and are essential for the infective cycle.

## Results

### Description of the ViroTreat algorithm

ViroTreat attempts to repurpose drugs as potential novel antiviral agents, based on their ability to invert the activity of VIPER-inferred MR protein modules controlling pro-infection programs. More specifically, the algorithm matches the set of virus-induced MRs that mechanistically regulate pro-infective programs with the context-specific Mechanism of Action (MoA) of a large repertoire of drugs, as inferred by VIPER-based analysis of RNA-seq profiles of drug perturbations vs. vehicle control (Fig. 1).

**Fig. 1.**
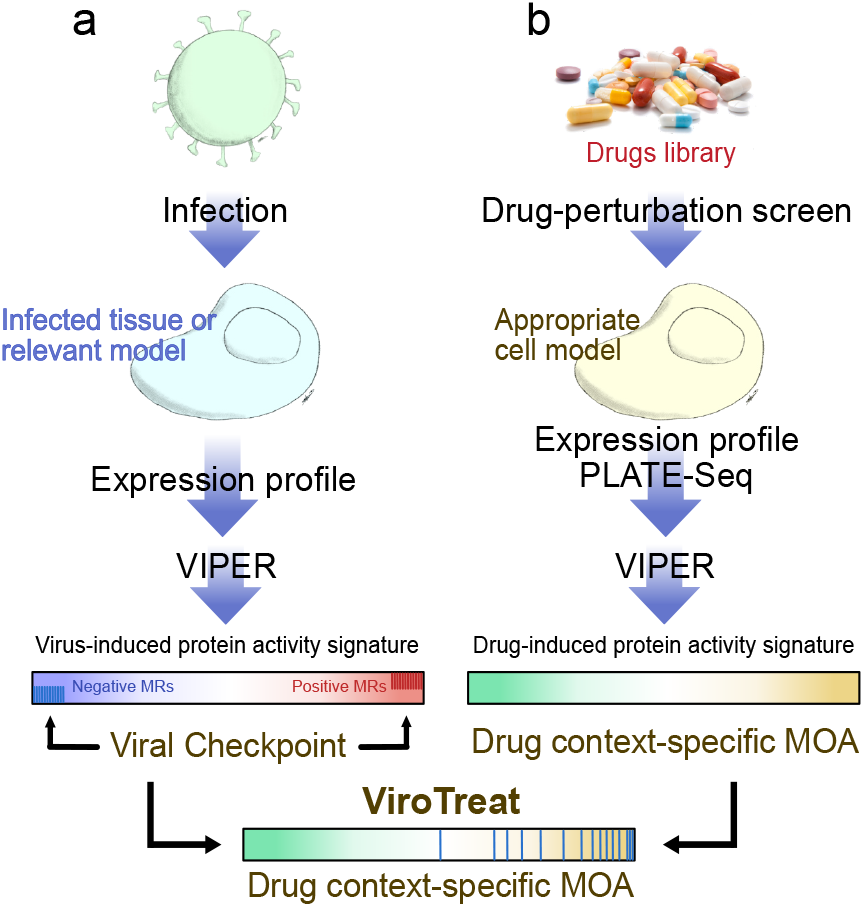
Schematic representation of the ViroTreat algorithm. **a**. Virus-induced MR proteins—the Viral Checkpoint—dissected by VIPER analysis of a gene expression signature, obtained by comparing an infected tissue or relevant model with non-infected mock controls. **b**. Context-specific drug MoA database, generated by perturbing an appropriate cell model with therapeutically relevant drug concentrations, followed by VIPER analysis of the drug-induced gene expression signatures to infer the drug-induced protein activity signature. ViroTreat prioritizes drugs able to activate the Viral Checkpoint’s negative MR proteins by quantifying the enrichment of such proteins on the drugs’ context-specific MoA.

First, MR proteins controlling regulatory programs induced by viral infection were identified by VIPER analysis comparing gene expression profiles from infected vs. non-infected (control) tissue (Fig. 1a). In cancer, we have shown that highly coordinated modules comprising between 10 and 50 proteins are necessary to control the state of a malignant cell(10, 19). We thus expect that an equally compact set of MR proteins would regulate the state of the host cells following SARS-CoV-2 infection and that a subset of these proteins would be responsible for implementing and sustaining cellular mechanisms critical for viral hijacking and replication. We thus focused on the top 50 most differentially active proteins following virus infection as “candidate SARS-CoV-2 infection MRs” (MRs hereafter, for simplicity) and refer to them, collectively, as the Viral Checkpoint (Fig. 1a).

Second, VIPER was also used to elucidate the MoA of 154 FDA-approved oncology drugs, where MoA is defined as the set of proteins differentially activated following drug perturbation of cell lines that recapitulate the regulatory network of the cellular population targeted by the virus(21). While this was done specifically in colon epithelial cells for this study, the analysis can be easily extended to assess drug MoA in other cellular contexts. Specifically, the RNA-seq profiles used in this analysis were generated at 24h (by Pooled Library Amplification for Transcriptome Expression (PLATE-Seq) assays(22)), following treatment of a colon adenocarcinoma cell line (LoVo) with a library of FDA-approved drugs and vehicle control (DMSO). To avoid assessing cell death or stress mechanisms, rather than drug MoA effects, drugs were titrated at their 48h IC_20_, as assessed by 10-point dose response curves (see methods for additional details). Resulting profiles were then used to assess the differential activity of regulatory proteins in drug vs. vehicle control-treated cells with the VIPER algorithm(12) (Fig. 1b).

Finally, drugs were prioritized based on their ability to invert the activity of SARS-CoV-2 Viral Checkpoint MRs, as assessed by enrichment analysis of the MRs in each drug context-specific MoA, using the analytic Rank Enrichment Analysis (aREA) algorithm(11, 12) (Fig. 1).

### SARS-CoV-2-induced MR signature

To elucidate MRs comprising the SARS-CoV-Viral Checkpoint, we analyzed publicly available single cell (scRNASeq) profiles of SARS-CoV-2 infected epithelial cells (Supplementary Table 1), including epithelial cell lines from both lung adenocarcinoma (Calu-3 and H1299)(23), and gastrointestinal organoid models from the ileum and colon(24). Single cell RNASeq analysis allows highly effective identification of individual virus-infected cells, which would otherwise represent only a minority of cells in culture. Moreover, single cell-based gene expression signatures—computed by comparing confirmed infected cells to non-infected controls—are less affected by contamination and dilution effects typical of bulk RNASeq signatures representing a mixture of infected and non-infected cells (see Supplementary Fig. 1 and Methods).

Single cell analysis revealed highly conserved differential protein activity signatures, as defined by the top 50 most differentially active candidate MRs (Viral Checkpoint), reflecting a highly conserved SARS-CoV-2 MR protein core, at each of the different time-points post-infection for which data was available from each model (*p <* 10^−40^, by 2-tailed aREA test, Fig. 2a and Supplementary Fig. 2a). Moreover, when comparing equivalent time-points, we observed significant conservation of the differentially active protein signature among lineage-related models (e.g., Calu-3 vs. H1299, at 12h, *p <* 10^−40^, Supplementary Fig. 2a).

**Fig. 2.**
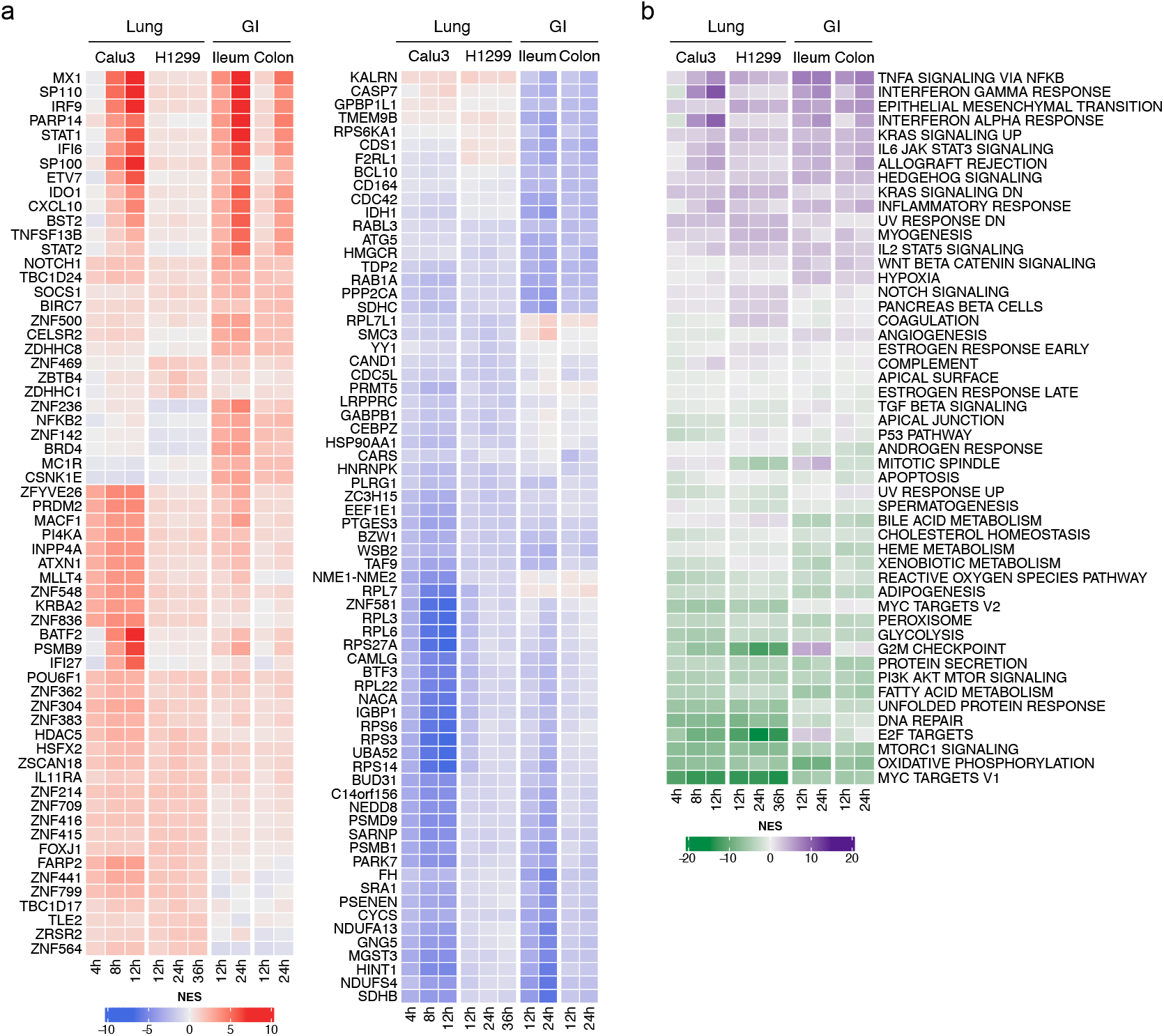
Changes in host cell protein activity in response to SARS-CoV-2 virus infection. **a**. Left, heatmap showing the activity of the top 10 most activated proteins in response to SARS-CoV-2 infection in each of the models and time-points profiled at the single-cell level. Right, heatmap showing the activity of the top 10 most inactivated proteins in response to SARS-CoV-2 infection in each of the models and time-points profiled at the single-cell level. **b**. Heatmap showing the enrichment of biological hallmarks in the SARS-CoV-2-induced protein activity signatures. Shown is the Normalized Enrichment Score (NES) estimated by the aREA algorithm, with purple color indicating enrichment in the over activated proteins and green color indicating enrichment in the inactivated proteins.

Moreover, such signature conservation was also observed among unrelated lineages, when comparing equivalent time-points (e.g., H1299 vs. colon non-transformed organoid at 24h, *p <* 0.01, Supplementary Fig. 2a). Taken together, these findings suggest the existence of a highly reproducible, SARS-CoV-2-mediated MR activity signature in epithelial cells, regardless of organ context (lung vs. gastrointestinal (GI)). Interestingly, however, inactivated MRs were significantly more conserved, both among models and between lineages, than activated MRs (*p <* 10^−6^, 2-tailed paired U-test, Supplementary Fig. 2b,c), suggesting a potentially distinct biological role for the activated vs. inactivated MR protein cores in SARS-CoV-2 infection.

The MR activity signatures detected by single cell analyses were also recapitulated by bulk-tissue analysis of SARS-CoV-2-infected epithelial cells (Supplementary Table 1), albeit at a slightly lower statistical significance, as we expected. These findings applied to both transformed models, including lung (Calu-3, H1299, and A549) and colon (Caco-2) adenocarcinoma, and normal human bronchial epithelial (NHBE) primary cells, as well as to more physiologic models, including lung organoids. As should be expected, MR conservation was more significant for models characterized by high infection rates (Supplementary Fig. 2a), likely due to signature dilution/contamination by a high proportion of non-infected cells in other models.

### MRs govern distinct biological functions

Enrichment analysis of SARS-CoV-2 MRs demonstrated a critical dichotomy of biological hallmark programs enriched in activated vs. inactivated MRs (Fig. 2b). Specifically, biological hallmarks enriched in activated MRs included inflammatory response, epithelial-to-mesenchymal transition (EMT) and interferon response. Indeed, among the top aberrantly activated MRs, we identified MX1, a protein induced by interferon I and II(25), the interferon regulator IRF9, and additional transcriptional regulators that mediate cellular response to interferons, such as STAT1 and STAT2(26) (Fig. 2a). Several poorly characterized, zing-finger proteins were also represented among the most differentially activated proteins, suggesting they may play a role in programs related to pathogen-mediated innate immunity.

In contrast, biological hallmarks enriched in inactivated MRs were strongly related to virus-mediated host-cell hijacking programs, such as PI3K signaling, unfolded protein response, DNA repair, and metabolic-related processes(27, 28) (Fig. reffig:fig2b). Consistent with this observation, the most significantly inactivated MRs included several ribosomal subunit members (such as RPS27A, RPS3, RPL3, RPS6, RPS14), as well as proteins involved in cell cycle arrest (UBA52)(29), translational regulation, and cellular metabolism (GABPB1)(30) (Fig. 2a).

### VIPER identifies key SARS-CoV-2-interacting proteins

To assess whether activated vs. inactivated MRs may represent a more effective target for drug-mediated reversal, we proceeded to assess whether either aberrantly activated or inactivated Viral Checkpoint MRs were enriched in host proteins already established as cognate binding partners of SARS-CoV-2 proteins. For this analysis, we leveraged a collection of 332 host proteins previously reported to be involved in protein-protein interactions (PPIs) with 26 of the 29 proteins encoded by the SARS-CoV-2 genome, as determined by mass-spec analysis of pull-down assays(5). Of these interactions, 90 were with proteins included in the 5,734 we analyzed by VIPER. Gene-Set Enrichment Analysis (GSEA)(31) revealed statistically significant enrichment of these 90 proteins in SARS-CoV-2 inactivated but not activated MRs, across all the evaluated single-cell protein activity signatures (*p <* 0.001, 2-tailed GSEA, Supplementary Fig. 3). This suggests that host cell proteins that physically interact with SARS-CoV-2 proteins are mostly inactivated in response to the infection.

Taken together, these results suggest an over-representation of inactivated MRs among host cell proteins that could critically modulate SARS-CoV-2 infectivity, thereby identifying the Viral Checkpoint module comprised of inactivated MRs as a candidate target and therapeutically actionable viral infection dependency.

### CRISPR assays confirm Viral Checkpoint MRs

To further confirm the biofunctional duality of Viral Checkpoint MRs, we also assessed their enrichment in proteins essential to the virus infectious cycle. Specifically, we evaluated their enrichment in proteins identified by functional CRISPR screens from two different studies, including using SARS-CoV-2 infected Vero(9) and Huh-7.5(7) cells. Consistent with our original observation and definition of the Viral Checkpoint signature, the 50 most inactivated candidate MRs—as determined by integrating results from both lung and GI models—were significantly enriched in antiviral genes, as assessed by each CRISPR screen (*p* < 10^−4^ and *p* < 10^−3^, respectively), as well as by their integration (Supplementary Fig. 4a–c, *p* < 10^−4^), while the 50 most activated MRs were not enriched in pro-viral genes (Supplementary Fig. 4d–f), confirming the requirement of Viral Checkpoint MRs inactivation for SARS-CoV-2 infectivity. These results however, could not be considered for the development of ViroTreat, since they became available after the candidate drugs in our study had already been prioritized and scheduled for validation.

### ViroTreat prioritization of FDA-approved drugs

We have shown that drug MoA—as represented by differentially active proteins in response to the drug and measured by VIPER analysis of drug perturbation profiles in lineage-matched cells—is well recapitulated *in vivo* and in explants when the activity of their MR proteins is conserved(21, 32). Among the 25 cell lines for which perturbational profiles had been generated in the PANACEA database (PANcancer Analysis of Chemical Entity Activity)(33), 2 cell lines, LoVo and NCI-H1973, are the closest lineage-matched representative models of the GI epithelial and lung epithelial cells, respectively. However, while LoVo (human colon cell line) showed significant conservation of MR proteins with a colon adenocarcinoma cell line susceptible to SARS-CoV-2 infection (Caco-2(34), Supplementary Fig. 5a,b), we did not observe MR protein activity conservation between NCI-H1793 cells and any of the three lung cell lines susceptible to SARS-CoV-2 infection (Calu-3, ACE2-A549 and H1299, Supplementary Fig. 5c–h). Based on these results and considering the availability of a compatible cell line model as a critical requirement to experimentally assess ViroTreat predicted drugs, we focused our drug repurposing efforts on the GI context.

We ranked drugs based on the enrichment of aberrantly inactivated MRs—following SARS-CoV-2 infection of GI cell lines and organoids—in proteins activated by each drug (ViroTreat algorithm, Fig. 1), based on the results of their perturbational assay. ViroTreat quantitative estimations were averaged across GI organoid models and evaluated time points. Among the 154 FDA-approved drugs profiled in LoVo cells, ViroTreat prioritized 22 (13 orally available and 9 intra-venous) at a highly conservative statistical threshold (*p <* 10^−5^, Bonferroni’s Corrected (BC)), see Fig. 3 and Supplementary Table 2).

**Fig. 3.**
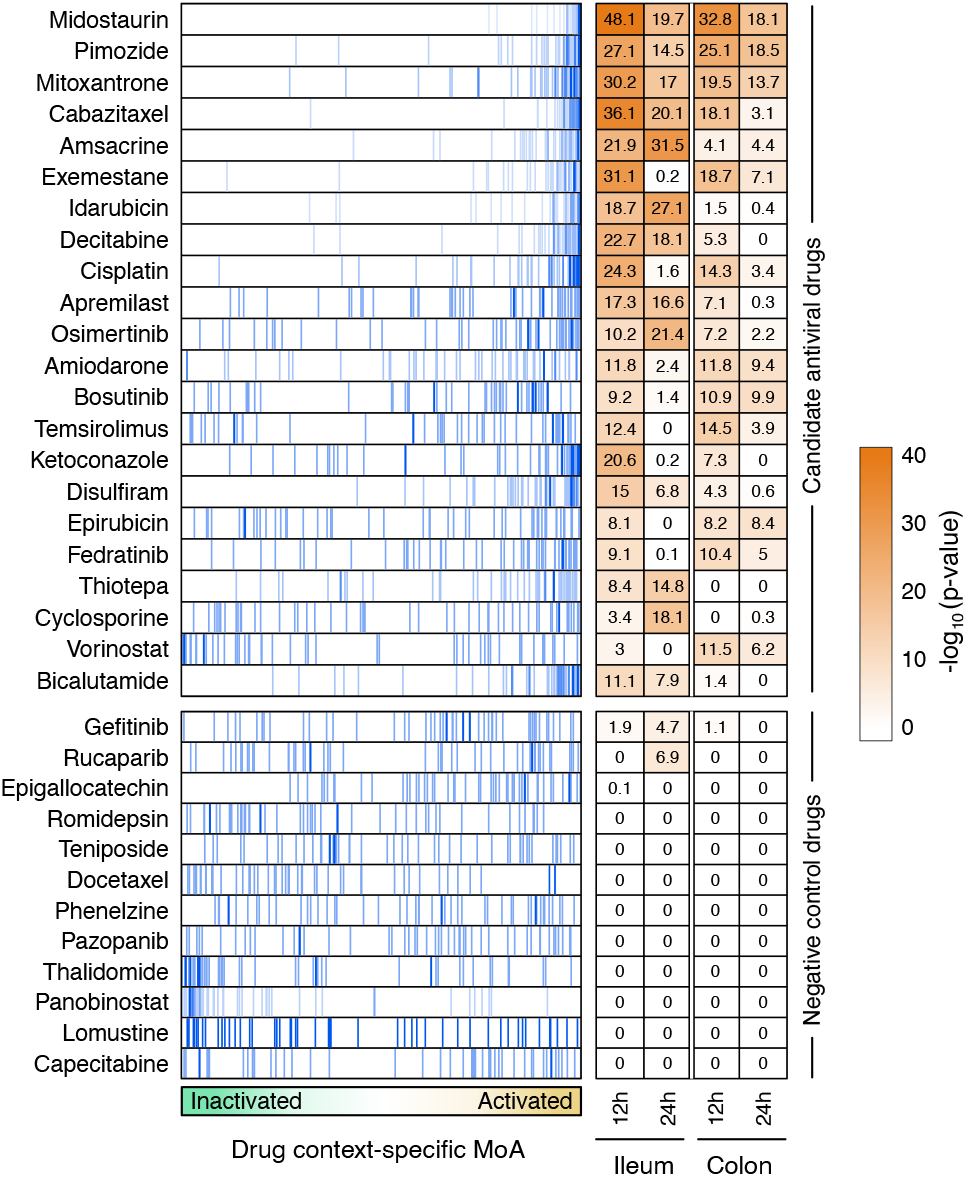
ViroTreat results for the GI models. Shown are the enrichment plot for the top 50 most inactivated (blue vertical lines) proteins, in response to SARS-CoV-2 infection (the negative component of the viral Checkpoint) of the ileum organoid for 12h, on the protein activity signature induced by the drug perturbations—drug context-specific MoA, represented by the green-orange color scale in the x-axis—of LoVo colon adenocarcinoma cells. The heatmap shows the Bonferroni’s corrected-log_10_ (p-value) estimated by ViroTreat. Shown are all the 22 candidate drugs (ViroTreat *p <* 10^−5^) and 12 drugs selected as negative controls (ViroTreat *p >* 0.01) in both ileum and colon-derived organoids at 12 and 24 hours post-infection.

### ViroTreat drugs inhibit SARS-CoV-2 infection

To provide proof-of-concept validation for the ViroTreat predictions, we first assessed drug-mediated inhibition of SARS-CoV-2 infection by ViroTreat-predicted vs. control drugs in the colon adenocarcinoma cell line (Caco-2) known to support SARS-CoV-2 infection(34).

For this assay, we considered all 13 ViroTreat-inferred orally-available drugs, as a more clinically relevant group, and the top 5 most significant intravenous (IV) drugs. As candidate negative controls, we selected 12 drugs—including 8 orally available agents and 4 IV drugs—not inferred as statistically significant by ViroTreat (*p* ≥0.01, Fig. 3 and Supplementary Table 2). Caco-2 cells were pre-treated for 24h prior to SARS-CoV-2 infection. Drug concentration was maintained through the entire infection time course and the relative infection levels and cell viability were assessed by immunofluorescence staining 24h post-infection (see methods and Fig. 4a). For each drug, the viability-normalized effect on SARS-CoV-2 infectivity (antiviral effect) was quantified as the log-ratio between infectivity and cell viability reduction relative to vehicle-treated (DMSO) controls (Supplementary Fig. 6). Since multiple concentrations were tested, the lowest concentration corresponding to a significant antiviral effect was reported (Supplementary Table 2). As a proof-of-concept for the ability of the methodology to identify drugs capable of reducing infectivity of SARS-CoV-2, we considered drugs to be validated only if their antiviral effect was statistically significant (False Discovery Rate (FDR) *<* 0.05) and they induced a decrease in infectivity of at least 20%.

**Fig. 4.**
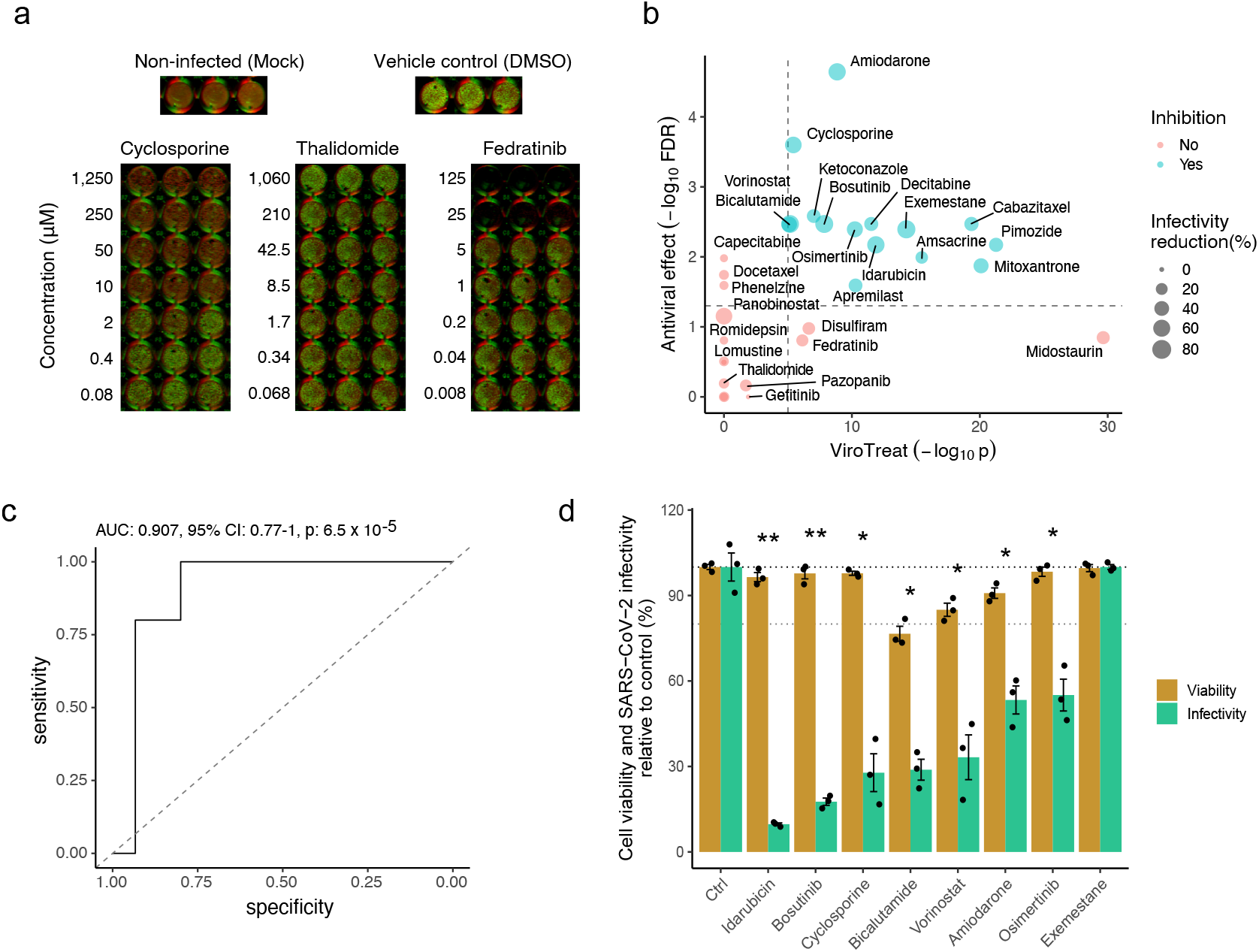
Experimental validation of ViroTreat predictions. **a**. Representative immunofluorescence images of non-infected (Mock) Caco-2 cells, vehicle control (DMSO) treated and SARS-CoV-2 infected cells, and representative examples of a drug showing significant antiviral effect (Cyclosporine), of a drug showing non-significant antiviral effect (Thalidomide) and a drug showing non-significant antiviral effect and cell toxicity (Fedratinib). Drug concentration (*µ*M) is indicated to the left of the images showing triplicated experiments. Cells were stained with DNA dye Draq5 (red) and anti-dsRNA antibody (green). **b**. Scatterplot showing the ViroTreat results (x-axis) compared to the specific antiviral effect (y-axis) experimentally evaluated in Caco-2 colon adenocarcinoma cells. The vertical and horizontal dashed lines represent the thresholds for statistical significance for ViroTreat (*p* = 10^−5^, BC) and specific anti-viral effect (FDR = 0.05), respectively. **c**. ROC analysis for the ViroTreat predictions, considering as positive response a specific antiviral effect at *FDR <* 0.05 with at least 20% reduction in infectivity. Estimated AUC, 95% CI and p-value are indicated in the plots. **d**. Effect of 8 drugs, showing the strongest reduction in SARS-CoV-2 infectivity in Caco-2 cells, on cell viability and SARS-CoV-2 infectivity of GI organoid-derived 2D primary cell cultures. Bars indicate the mean *±* Standard Error of the Mean (SEM). Antiviral effect: * FDR *<* 0.05, ** FDR *<* 0.01.

Of 18 drugs predicted to inhibit SARS-CoV-2 infection, 15 (83%) showed statistically significant antiviral effect. In contrast, none of the 12 drugs selected as potential negative controls showed significant antiviral effect (Fig. 4b and Supplementary Table 2), demonstrating a significant enrichment of ViroTreat results in drugs with antiviral activity (*p <* 10^−5^, 1-tailed Fisher’s exact test (FET)). Consistently, the Receiver Operating Characteristic (ROC) had an Area Under the Curve (AUC) = 0.907 (95% confidence interval (CI): 0.77– 0.91), which is highly statistically significant (*p <* 10^−4^, Fig. 4c), demonstrating the predictive power of ViroTreat in this proof-of-concept.

To further assess the pathogen-specific nature of ViroTreat predictions, we tested the ability of the 8 ViroTreat-inferred drugs showing the strongest inhibition of SARS-CoV-2 infectivity, to inhibit rotavirus infection and replication in Caco-2 cells. Interestingly, none of these drugs significantly impaired rotavirus infectivity (Supplementary Fig. 7 and Supplementary Table 2), showing that ViroTreat-inferred antiviral effects cannot be attributed to generalized impairment of host cellular functions universally required for viral infection, but rather to activation of MRs specific to SARS-CoV-2 infection.

To also assess whether the antiviral activity of ViroTreat-predicted oncology drugs in Caco-2 cells might possibly be attributed to their antineoplastic effects in a cancer cell context, we evaluated the antiviral properties of the top 8 drugs in non-transformed, human GI organoid-derived 2D primary cell cultures. When tested on this more physiologic context, 7 of the 8 assayed drugs, including idarubicin, bosutinib, cyclosporine, bicalutamide, vorinostat, amiodarone and osimertinib, demonstrated significant antiviral effect against SARS-CoV-2 based on our original criteria (FDR *<* 0.05 and decrease in SARS-CoV-2 infectivity of at least 20%, Fig. 4d and Supplementary Fig. 7). Except for bicalutamide, which exerted its antiviral effect at a 125-fold higher concentration, all drugs were tested at concentrations comparable to their 48h IC_20_ in LoVo cells, representing the highest sub-toxic concentration usable for optimal MoA elucidation. These findings suggest that ViroTreat can apply the molecular characterization of a drug’s MoA, as obtained by the measured effect of the drug on protein activity levels in tissue lineage-matched, neoplastic cell line models, to prioritize and repurpose drugs with potential antiviral activity in both infected tumor models as well as non-transformed human organoid-derived 2D primary cell cultures.

Finally, to test the tissue lineage context-specificity of ViroTreat predictions, we assessed the antiviral effect of the 8 ViroTreat predicted drugs for the GI context showing the strongest inhibition of SARS-CoV-2 infection in Caco-2, in lung adenocarcinoma cell line models (Calu-3 and ACE2-A549). Interestingly, only cyclosporine and osimertinib showed a significant antiviral effect (FDR *<* 0.05 and ≥ 20% infectivity decrease), while amiodarone, apremilast, bicalutamide, bosutinib, exemestane, and pimozide did not (Supplementary Fig. 8 and Supplementary Table 2). These results highlight the relevance of lineage context-specificity when prioritizing drugs with ViroTreat.

## Discussion

While immunization remains the principal strategy to mitigate SARS-CoV-2 transmission and its associated clinical morbidity, a significant fraction of the population is likely to remain unvaccinated. In addition, protection against COVID-19 is imperfect and incremental resistance to current vaccines, in the form of more transmissible and deadly SARS-CoV-2 variants, has already been documented(8, 35, 36). As a result, identification of safe and effective therapies for COVID-19 patients—especially via short course, non-toxic, and orally administered drugs—remains a major priority to address for the current pandemic. This therapeutic strategy would be especially valuable for patients with mild-to-moderate symptoms in whom hospitalization may be prevented. Since accurate medical devices for assessing blood oxygen saturation have been cleared for in-home use, treating patients early in the disease could change the pandemic landscape by allowing treatment in an outpatient context.

To provide a proof-of-concept for systematically addressing these kinds of challenges and unmet needs, we developed ViroTreat as a mechanism-based framework for repurposing drugs, based on their ability to reprogram host cells to a state refractory to virus hijacking. In contrast to previous host-centric approaches aimed at targeting single host cell proteins that directly interact with the viral proteome, ViroTreat was designed to target an entire set of MR proteins, whose concerted regulatory activity is responsible for implementing and maintaining a virus replication-permissive state in host cells, as elucidated by VIPER-based Master Regulator analysis of infected vs. control cells. By doing so, ViroTreat expands the one disease/one target/one drug paradigm to one centered on reversing the activity of an entire protein module (i.e, Viral Checkpoint) based on the accurate assessment of each drug’s proteome-wide MoA, as dissected from perturbational profile data. Such a holistic approach to matching disease dependencies to drug MoA overcomes the inherent limitations of drug repurposing efforts that focus on inhibitors of individual proteins or single pathways to thwart viral infectivity as part of a host cell-directed strategy.

Elucidation of Viral Checkpoint MRs requires availability of gene expression signatures that accurately reflect virus-mediated changes in the host cell transcriptome. Thus, to avoid confounding effects by model-idiosyncratic mechanisms and to ensure identification of more universal and reproducible MR proteins, we dissected the Viral Check-point from multiple, complementary models, including transformed cell lines and normal 3D-organoid cultures representing both airway and GI epithelium lineages. In addition, to avoid additional confounding effects arising from infection heterogeneity, we performed VIPER analysis at the single cell level, thereby mitigating the contribution of non-infected cells, which represent the majority of the tissue, based on reads mapping to the SARS-CoV-2 genome. Similarly, we avoided confounding effects arising from single cell transcriptional state heterogeneity by comparing each infected cell to a small pool of the closest non-infected cells, based on MR analysis, as controls. Finally, to achieve context-specific understanding of drug MoA, the analysis was performed in tissues reflective of the biology of infected cells based on conservation of their most differentially active MRs, as previously described(21, 32).

The ViroTreat framework prioritizes drugs from a predefined library used to generate perturbational assays. For this proof-of-concept, we maximized the translational potential of drug predictions, by focusing our analysis on FDA-approved drugs used primarily in an oncology setting; with particular emphasis on orally available drugs. However, the approach can be easily extended to explore a much larger library of pharmacological compounds. Moreover, the database of drug context-specific MoA can be generated independently and prior to the identification, isolation and characterization of a viral pathogen of interest, making it readily available for current as well as future pandemics.

In addition, while most studies have focused on drugs that act as high affinity inhibitors of target proteins(5–7, 9, 37–39), to our knowledge, this is the first study to focus on pharmacologic agents predicted to activate rather than inhibit an entire module of Master Regulator proteins whose inactivation by the virus was found necessary for its replication cycle. By inducing drug-mediated reversion of Viral Checkpoint activity, we successfully reprogrammed host cells to a state of “viral regulatory contraception”, thereby significantly compromising the ability of the virus to hijack host cell machinery required for its infective cycle. Accordingly, ViroTreat allows identification of a global, pathway-independent, therapeutically actionable set of pharmacological agents acting at multiple, key nodes of the host cell’s regulatory network, and as such, most likely generalizable across a broad spectrum of current and yet-to-evolve variants of SARS-CoV-2. Notably, the validation rate for predicted drugs in independent assays was extremely high. And, importantly, in a completely unbiased fashion, Virotreat predicted antiviral activity against SARS-CoV-2 of drugs that have recently emerged as potential host cell-targeting antivirals, among them, cyclosporine(40), amiodarone(41), pimozide(42), mitoxantrone(43), osimertinib(44), bosutinib(45), and bicalutamide(46). Moreover, three of the Virotreat-predicted drugs—cyclosporine (NCT04492891), amiodarone (NCT04351763), and bicalutamide (NCT04509999)—are being evaluated in clinical trials for their safety and efficacy in persons with SARS-CoV-2 infection.

Among the methodological limitations, the most critical one is the need to obtain physiologic models to identify appropriate infection signatures, generate relevant drug perturbational profiles, and validate predicted drugs. In addition, there are also challenges in assessing the optimal concentration at which each compound should be profiled.

From a translational perspective, in the setting of both the current and future pandemics, as well as for recurrent epidemics such as those caused by influenza and other viral pathogens, the Viral Checkpoint framework can leverage bulk and single-cell profiles from infected cells to quickly identify the precise set of MR proteins responsible for creating a virus infection-friendly environment in the host cell. Once identified, independent of the specific viral pathogen, potential therapeutic agents can be efficiently prioritized by ViroTreat, using readily available—and relatively inexpensive—perturbational databases to elucidate context-specific, proteome-wide drug MoA. Host cell-directed therapies shown to be effective in cell line and organoid models based on such predictions can then undergo rapid validation in more physiologic contexts, prior to testing in human trials designed to evaluate their safety and therapeutic value in the clinical setting.

## ACKNOWLEDGEMENTS

We would like to thank Dr. Vibor Laketa and Dr. Sylvia Olberg for their support through the Infectious Diseases Imaging Platform (IDIP) and Tatiana Alvarez for original artwork. This research was supported by the following NIH grants to A.C.: R35 CA197745 (Outstanding Investigator Award); U01 CA217858 (Cancer Target Discovery and Development); S10 OD012351 and S10 OD021764 (Shared Instrument Grants); by grants to S.B.: Deutsche Forschungsgemeinschaft (DFG) project numbers 415089553 (Heisenberg program), 240245660 (SFB1129), 278001972 (TRR186), and 272983813 (TRR179), and from the state of Baden Wuerttem-berg (AZ: 33.7533.-6-21/5/1) and the Bundesministerium Bildung und Forschung (BMBF) (01KI20198A) and within the Network University Medicine - Organo-Strat COVID-19; by grants to M.L.S.: BMBF (01KI20239B) and DFG project 416072091; by grant to T.A.: ERC Consolidator grant METACELL (grant number 773089); by grant to M.K.J.: BCPM grant to Thomas Geiser (Department of Pulmonary Medicine, University Hospital/DBMR), and support from the Department of Urology of the Bern University Hospital; and to M.J.A.: research support from Karyopharm Therapeutics, Inc.

## Author Contributions

P.L., G.B., M.K.J., S.B., A.C., and M.J.A. conceived this work. M.L.S., S.T., P.D., T.A., F.L.M, and M.D.M performed experiments. R.B.R. and S.P. performed experiments and generated the drug perturbational data. P.L., X.S., and M.J.A. performed analysis. G.B., C.K., T.A., M.K.J, A.C., S.B., and M.J.A. supervised experiments and data analysis. P.L., M.L.S., G.B., A.C., S.B., and M.J.A. wrote the manuscript. P.L., M.L.S., G.B., M.K.J., A.C., S.B., and M.J.A. reviewed the manuscript. All authors approved the final manuscript.

## Methods

### Cell lines

Vero E6 (ATCC CRL-1586) and Caco-2 (ATCC HTB-37) cells were maintained in DMEM supplemented with 10% fetal bovine serum and 1% penicillin/streptomycin.

### GI organoids

Human tissue was received from colon resection from the University Hospital Heidelberg. This study was carried out in accordance with the recommendations of the University Hospital Heidelberg with informed written consent from all subjects in accordance with the Declaration of Helsinki. All samples were received and maintained in an anonymized manner. The protocol was approved by the “Ethics commission of the University Hospital Heidelberg” under the protocol S-443/2017. Stem cells containing crypts were isolated following previously described protocols(47). Organoids were passaged and maintained in basal and differentiation culture media (Supplementary Table 3) as previously described(47).

### Viruses

SARS-CoV-2 (strain BavPat1) was obtained from the European Virology Archive. The virus was amplified in Vero E6 cells and used at a passage 3 for all experiments as previously described(24, 34).

### SARS-CoV-2 infection assay

20,000 cells were seeded per well into a 96-well dish 24 hours prior to drug treatment. 100 *µ*L of media containing the highest drug concentration was added to the first well. Six serial 1:5 dilutions were made (all samples were performed in triplicate). Drugs were incubated on cells for 24 hours. Prior to infections, fresh drugs were replaced and SARS-CoV-2 at Multiplicity Of Infection (MOI) 3 was added to each well. 24 hours post-infection cells were fixed in 4% paraformaldehyde (PFA) for 10 mins at Room Temperature (RT). PFA was removed and cells were washed twice in 1X PBS and then permeabilized for 10 mins at RT in 0.5% Triton-X. Cells were blocked in a 1:2 dilution of Li-Cor blocking buffer (Li-Cor) for 30 mins at RT. Cells were stained with 1/1000 dilution anti-dsRNA (J2, SCIONS) for 1h at RT as marker of infected cells as previously described(34). Cells were washed three times with 0.1% Tween in PBS. Secondary antibody goat antimouse IR 800 (Thermo) and DNA dye Draq5 (Thermo) were diluted 1/10,000 in blocking buffer and incubated for 1h at RT. Cells were washed three times with 0.1% Tween/PBS. Cells were imaged in 1X PBS on a LICOR imager. Effect of drugs were analyzed by comparing the average fluorescence of mock treated cells to drug treated cells. Draq5 staining was used to determined cell viability.

### Rotavirus infection assay

40,000 cells were seeded per well into a collagen-coated 96-well dish 24 hours prior to drug treatment. 100 *µ*L of media containing the highest drug concentration was added to the first well. Six serial 1:5 dilutions were made (all samples were performed in triplicate). Drugs were incubated on cells for 24 hours. Media was removed and cells were washed 2X with serum-free media and were infected with WT SA11 Rotavirus expressing mKate at MOI 0.1 (calculated in MA104 cells) diluted in serum-free media. Rotavirus was previously activated for 30 minutes at 37°C in serum-free media containing 2 *µ*g/ml trypsin. Infection was allowed to proceed for 1 hour. Following infection, virus was removed and cells were washed 1X with serum-free media. Media containing drugs and 0.5 *µ*g/ml trypsin were added back to cells to allow for Rotavirus propagation. 24 hours post-infection cells were fixed with 2% PFA for 15 mins and then stained with DAPI. Cells were imaged in 1X PBS on a Cell Discoverer 7 using a 5X objective. Quantifications of infection was carried out by quantifying the number of infected cells (mKate positive cells) in infected and not infected samples using CellProfiler.

### SARS-CoV-2 infection of human colon organoids-derived 2D primary cell cultures

Organoids were cultured in 24-well plates in basal medium for 5–7 days following the original protocol of Sato and co-workers(47). To obtain human colon organoids-derived 2D primary cell cultures, the medium was removed from the 24-well plates, organoids were washed 1X with cold PBS and spun (450g for 5 mins). PBS was removed and organoids were digested with 0.5% Trypsin-EDTA (Life technologies) for 5 mins at 37°C. Digestion was stopped by addition of serum containing medium. Digested-organoids were spun again at 450g for 5 mins and the supernatant was removed and digested organoids were re-suspended in basal media at a ratio of 250 *µ*L media/well (corresponding to approximately 400 organoids per ml). Prior seeding, the 48-well tissue culture plates were coated with 2.5% human collagen in water for 1 h at 37°C. The collagen mixture was removed from the 48-well plate and 250 *µ*L of trypsin-digested organoids (corresponding to about 100 digested organoids) were added to each well. 48 hours post-seeding differentiation media (Supplementary Table 3) was added to cells and 4 days post-differentiation cells were treated with drugs at the indicated concentrations for 2 hours prior to SARS-CoV-2 infection. Media containing drugs was removed and 10^6^ Focus Forming Units (FFU) (as determined in Vero cells) of SARS-CoV-2 was added to each well for 1 hour at 37°C. Following 1 hour incubation, virus was removed and fresh differentiation media containing drugs was added to cells. 24 hours post-infection RNA was harvested, and virus replication was monitored by RT-qPCR.

### Estimation of the antiviral effect

We define the antiviral effect of a drug as its viability-normalized effect on SARS-CoV-2 infectivity. The antiviral effect was quantified as the log-ratio between infectivity and cell viability reduction relative to vehicle-treated controls. Statistical significance was estimated by Student’s t-test for each evaluated drug concentration, and multiple-hypothesis testing due to the multiple evaluated concentrations was corrected using the conservative Bonferroni’s method. Multiple hypothesis testing due to multiple evaluated drugs was further corrected by Benjamini-Hochberg False Discovery Rate (FDR).

### RNA isolation, cDNA, and RT-qPCR

RNA was harvested from cells using RNAeasy RNA extraction kit (Qiagen) as per manufactures instructions. cDNA was made using iSCRIPT reverse transcriptase (BioRad) from 250 ng of total RNA as per manufactures instructions. RT-qPCR was performed using iTaq SYBR green (Bio-Rad) as per manufacturer’s instructions. TBP or HPRT1 were used as normalizing genes. See Supplementary Table 4 for primers used.

### VIPER analysis of bulk RNA-Seq datasets

The source for all the datasets is listed in Supplementary Table 1. RNA-Seq raw-counts data for Calu-3, H1299 and Caco-2 cell line models were obtained from Gene Expression Omnibus Database (Gene Expression Omnibus (GEO), GSE148729)(23). Raw-counts data for A549 cell line, Normal Human Bronchial Epithelial (NHBE) primary cells, a post-mortem lung tissue sample from a COVID-19 patient and a healthy human lung biopsy were downloaded from GEO (GSE147507)(48). Normalized data (Transcript per Kilobase Million, TPM) for lung organoids were downloaded from GEO (GSE160435). Raw-count data was normalized using the variance stabilization transformation (VST) procedure as implemented in the DESeq package from Bioconductor(49).

Differential gene expression signatures for the Wyler’s dataset(23) (GSE148729) were computed by comparing the SARS-CoV-2 infected samples against the centroid—i.e. the average expression of each gene—of the closest matched non-infected (mock) samples as identified by unsupervised clustering. Specifically, we first performed K-means cluster analysis of the normalized gene expression profiles. The optimal number of clusters was estimated by silhouette-score analysis as implemented in the “fviz_nbclust” function of the “factoextra” package (https://cran.r-project.org/web/packages/factoextra/index.html). Cluster solutions were evaluated from *k* = 2 to *k* = 10 and the solution with the highest average of silhouette scores was considered as optimal. Based on the optimal cluster solution, we selected as reference for each infected sample the centroid of the mock samples within the same cluster. In cases of clusters constituted by infected samples only, the centroid of the mock samples in the closest cluster were used as reference. Because a two clusters solution was estimated as optimal for all cluster analysis, the other cluster was the trivial closest cluster solution in all cases. Cluster solutions with less than two samples per cluster were considered ineffective. For Calu-3 cell line, we noticed that samples associated to the two series (series-1 and series-2) clustered separately—i.e. samples clustered according to series memberships. To avoid possible batch effects in the analysis, the samples of these two series were re-clustered separately to identify the best matched mock control samples in each series independently. For series-1, the mock samples at 4h and 24h clustered together and were used as reference to compute the differential expression signatures of all the Calu-3 SARS-CoV-2 infected samples. For series-2, three mock samples, including one mock sample at 4h and two mock samples at 12h clustered together and were used as reference to compute the differential expression signatures for all the Calu-3 SARS-CoV-2 infected samples. Of note, in series-2, one mock sample at 4h (GSM4477923) clustered separately from all the other samples with a silhouette score of zero which indicates no clear cluster assignment. This sample was considered as outlier and excluded from the downstream analysis. For the Caco-2 cell line, the centroid of the 4h mock samples was used as reference to compute the differential expression signatures of the SARS-CoV-2 infected samples at 4h and 12h, while the centroid of 24h mock samples was used as reference to compute the differential expression signatures of the 24h SARS-CoV-2 infected samples. For the H1299 cell line, the centroid of the 4h mock samples was used as reference to compute the differential expression signatures of the SARS-CoV-2 infected samples at 4h and 12h; and the centroid of the 36h mock samples was used as reference to compute the differential expression signatures of the 36h SARS-CoV-2 infected samples.

Differential gene expression signatures for the Blanco-Melo’s dataset(48) (GSE147507) were computed using the centroid of the matched—i.e. same cell line or primary cells—mock control samples as reference. For the post-mortem human lung sample from a COVID-19 patient, the differential gene expression signature was computed using the healthy human lung biopsy samples as reference.

Differential gene expression signatures for the lung organoid sample was computed using as reference its matched mock control sample.

The differential activity of 5,734 proteins, including 1,723 transcription factors, 630 co-transcription factors, and 3,381 signaling proteins, was estimated for each of the differential gene expression signatures with the VIPER algorithm(12), using matched context-specific models of transcriptional regulation. Lung, colon and rectal adenocarcinoma context-specific models of transcriptional regulation were reverse-engineered, based on 517 lung, 459 colon and 167 rectal adenocarcinoma samples in The Cancer Genome Atlas (TCGA) with the ARACNe algorithm(13, 50), as discussed in(19). While, ideally, regulatory networks from non-cancer-related epithelial cells may have been more appropriate, use of cancer-related regulatory networks is justified by the high conservation of protein transcriptional targets in cancer-related and normal cells from the same lineage(14). The regulatory models are available as part of the arcane.networks R package from Bioconductor (https://www.bioconductor.org/packages/release/data/experiment/html/aracne.networks.html). Specifically, protein activity signatures in response to SARS-CoV-2 infection of the lung adenocarcinoma cell lines (Calu-3, H1299 and A549), lung organoids and human lung tissue samples were inferred with the VIPER algorithm using the lung adenocarcinoma context-specific network. Protein activity signatures for Caco-2 colorectal carcinoma cell line were estimate with the metaVIPER algorithm(16) using the colon and rectal adenocarcinoma context-specific networks.

The VIPER-inferred protein activity signatures of infected samples at the same time point in the same cell line were integrated using the Stouffer method(51).

### VIPER analysis of scRNA-Seq datasets

Single-cell (sc)RNAseq count matrices, based on Unique Molecular Identifiers (UMIs), for Calu-3 and H1299 lung adenocarcinoma cell lines were downloaded from GEO (GSE148729). Both count matrices were already filtered for low quality cells as described(23). Count matrices (UMI) from ileum and colon organoids were made available by Boulant lab and are also publicly available on GEO (GSE156760). Count matrices were filtered for low quality cells as described by Triana et al.(24).

In contrast to bulk RNASeq profiles, single cell RNASeq profiles (scRNASeq) allow effective identification of the individual cells likely to be infected by the virus, which commonly represent a minority of cells in a culture. For this study, therefore, we defined cells to be infected if they present at least one sequenced read mapped to the SARS-CoV-2 genome. Critically, gene expression signatures based on scRNASeq profiles, as computed by comparing *bona fide* infected cells to non-infected controls, are less affected by contamination and dilution effects characteristic of bulk RNASeq-derived signatures, resulting from a variable proportion of infected vs. noninfected cells.

To account for confounding effects and gene expression profile heterogeneity associated with mechanisms that are independent of viral infection(23, 24)—such as cell cycle and the use of models derived from cancer cell lines(52)—differential expression signatures between infected and non-infected single cells were computed by comparing each infected cell to its *k* = 50 closest non-infected ones (Supplementary fig. 1). This approach significantly improved accuracy and reproducibility of differential gene expression signatures, including across different cell lines, by minimizing confounding effects not associated with viral infection. To identify mock controls cells for each individual infected cell we transformed the count matrices to count per million (CPM) and subsequently to VIPER-inferred protein activity signatures. Briefly, gene expression profiles were transformed to differential gene expression signatures using the “scale” method—i.e. z-score transformation—as implemented in the VIPER package(12). Then, using lung adenocarcinoma context-specific models of transcriptional regulation, we transformed the single-cell gene expression signature matrices for Calu-3 and H1299 cell lines to VIPER-inferred protein activity signature matrices. Similarly, using colon and rectal adenocarcinoma context-specific networks, we transformed the single-cell gene expression signature matrices for ileum and colon organoids to the corresponding metaVIPER-inferred protein activity signature matrices.

The phenotypic state similarity between cells of the same dataset was quantified by the euclidean distance, calculated based on the top 100 principal components of the VIPER-inferred protein activity matrix. Briefly, the Singular Value Decomposition (SVD) was used to estimate the matrix of cells by eigenproteins (principal components), and linear regression analysis was used to identify the components (eigenprotein vectors) significantly associated to the viral infection, expressed as the sum of the normalized UMI viral counts—counts mapping to the SARS-CoV-2 genome. For ileum and colon, the vectors of viral counts were generated by summing the normalized counts generated by targeted sequencing analysis(24). Principal components significantly associated with infection (*p <* 0.05) were removed from the Principal Component Analysis (PCA) space. Next, we performed a K-Nearest Neighbors (KNN) analysis in the dimensionally reduced PCA space, considering the top 100 infection-independent principal components, to identify the phenotypically closest 50 mock cells for each of the infected cells. The KNN analysis was performed using the FNN package(53). The 50 phenotypically closest mock cells were used as reference to compute the SARS-CoV-2-induced differential gene expression signature for each of the infected cells. Specifically, the differential gene expression signature for each infected cell was estimated by subtracting the mean expression of the 50 phenotypically closest mock cells and dividing by their standard deviation. For Calu-3 and H1299 cell lines, we considered as “SARS-CoV-2-infected” all the cells with at least 1 sequencing read mapping to the SARS-CoV-2 genome. For ileum and colon, we considered as “SARS-CoV-2-infected”, all cells identified by targeted sequencing(24).

The differential gene expression signatures of SARS-CoV-2 infected cells were transformed to inferred protein activity signatures by VIPER and metaVIPER algorithms, as described above.

Single-cell protein activity signatures of each data set were integrated by arithmetic mean at each available time point for each cell line.

### Similarity of VIPER-inferred protein activity signatures

The conservation of MR proteins between VIPER-inferred protein activity signatures was quantified by the reciprocal enrichment of the top 25 most activated, and the top 25 most inactivated proteins in signature *S*_1_ in proteins differentially active in signature *S*_2_ and vice versa(54), as implemented by the viperSimilarity() function in the viper package from Bioconductor.

### Enrichment of biological hallmarks on SARS-CoV-2 infection-induced protein activity signatures

Hallmarks gene sets (v.7.2) were downloaded from the molecular signatures database (MSigDB) website (http://www.gsea-msigdb.org/gsea/msigdb/collections.jsp). Enrichment of the MsigBD biological hallmarks protein-sets on the SARS-CoV-2 induced, VIPER-inferred protein activity signatures, with the aREA algorithm(12).

### Enrichment of Viral Checkpoint MRs on infection essential genes identified by CRISPR screens

CRISPR screen results (z-score) were downloaded from the supplementary data of Wei et.al(9) (Vero-E6 cells) and Schneider et.(7) (Huh-7.5 cells). Z-scores were integrated across all experimental conditions for each cell line using the Stouffer’s method(51). Enrichment of the top 50 most activated, and the top 50 most inactivated proteins in response to SARS-CoV-2 infection, obtained after integrating (average) all 10 single-cell protein activity signatures, on each CRISPR experiment z-score signature, and on their Stouffer’s integration, were estimated by GSEA. Normalized Enrichment Score (NES) and p-value were estimated by permuting the genes in the CRISPR signatures 10,000 times uniformly at random. SARS-CoV-2 inactivated MRs essential for infectivity were identified as the genes in the leading-edge for the GSEA of the inactivated MRs on the integrated CRISPR screen signature.

### Enrichment of SARS-CoV-2 interacting protein on host proteins differentially active in response to SARS-CoV-2 infection

A list of 332 SARS-CoV-2 interacting proteins was obtained from the supplementary materials of Gordon et al.(5). 90 of the 332 interacting proteins were represented among the regulatory proteins for which we could infer their activity. Enrichment analysis of this 90 SARS-CoV-2 interacting proteins on the VIPER-inferred protein activity signatures was performed by GSEA. NES and p-values were estimated by permuting the VIPER-inferred protein activity signatures 10,000 times uniformly at random.

### ViroTreat analysis

We have previously shown that tumor check-points can be pharmacologically switched, either off(11, 15, 20, 55, 56) or on(19), leading to their collapse and loss of viability or gain of associated functional properties, respectively. This observation was instrumental for the development and validation of the NY CLIA certified, VIPER-based methodology OncoTreat, for the prioritization of small molecule compounds that can either inactivate or activate a tumor checkpoint on a sample-by-sample basis, with critical applications in precision oncology(11). Based on the successful outcomes observed with OncoTreat when evaluated in the context of tumor suppression, we sought to develop a novel, analogous algorithm, ViroTreat, to identify small molecule compounds capable of suppressing viral infection by targeting the Viral Checkpoint module. Similar to its use in cancer, ViroTreat systematically assesses and prioritizes a small-molecule compound’s ability to reverse the activity of a set of MR proteins based on large-scale drug perturbation assays in cell lines that recapitulate (a) the regulatory model of the target cellular population and (b) the activity of MR proteins. Specifically, perturbational assay data are comprised of RNASeq profiles generated at 24h (by PLATE-Seq assays(22)), following treatment of MR-matched cell lines with a library of FDA-approved and late-stage experimental drugs (in Phase 2 and 3 clinical trials) and DMSO as control. These profiles are then used to assess the differential activity of relevant MRs in drug vs. DMSO-treated cells. Finally, enrichment of MR proteins in proteins whose activity has been inverted by the drug is computed by protein set enrichment analysis (PSEA) using the aREA algorithm(11, 57). The RNASeq profiles used for ViroTreat analysis were generated at 24h following treatment of LoVo cells with a repertoire of 154 FDA-approved oncology drugs. Perturbations were performed at each drug’s highest sublethal concentration (48h IC_20_) or maximum serum concentration (C_max_) at its Maximum Tolerated Dose (MTD), whichever was lower. This was done to prevent confounding effects, unrelated to the drug MoA, resulting from cell death or stress pathway activation. RNASeq data was generated using PLATE-Seq, a fully automated, 96-well based assay(22) (Supplementary Table 2).

### Code availability

All the code used in this work is freely available for research purposes. VIPER and aREA algorithms are part of the “viper” R-system’s package available from Bioconductor. The lung adenocarcinoma, colon and rectal context-specific interactomes are available as part of the “aracne.networks” R-system’s package from Bioconductor.

## Supplementary Figures and Tables

**Supplementary Figure 1.**
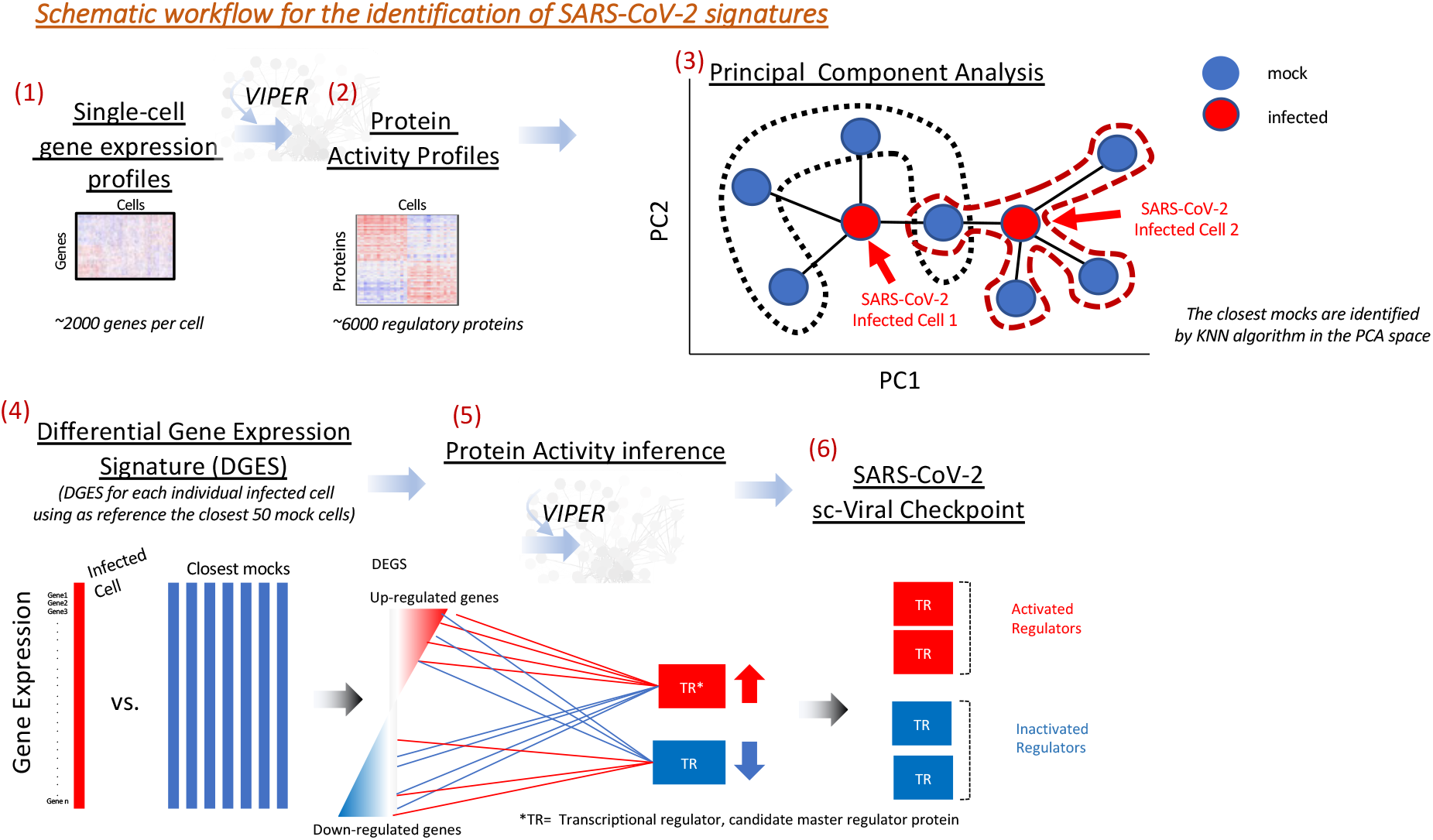
Diagram showing the workflow used to compute the protein activity signatures induced by SARS-CoV-2 infection from scRNA-Seq data. Normalized single-cell gene expression profiles for all cells of the same model (i.e. Calu3, H1299, colon and ileum) were transformed to differential gene expression signatures by applying the z-score procedure. Single-cell differential gene expression signatures were then transformed to protein activity profiles by applying the VIPER algorithm with context-specific regulatory networks. A Principal Component Analysis (PCA) was performed on these VIPER-inferred protein activity profiles. For each infected cell the closest 50 mock cells in the PCA space were selected as reference to compute a SARS-CoV-2 induced differential gene expression signature. The VIPER algorithm was then applied to these SARS-CoV-2 induced differential gene expression signatures to infer SARS-CoV-2 induced protein activity signatures.

**Supplementary Figure 2.**
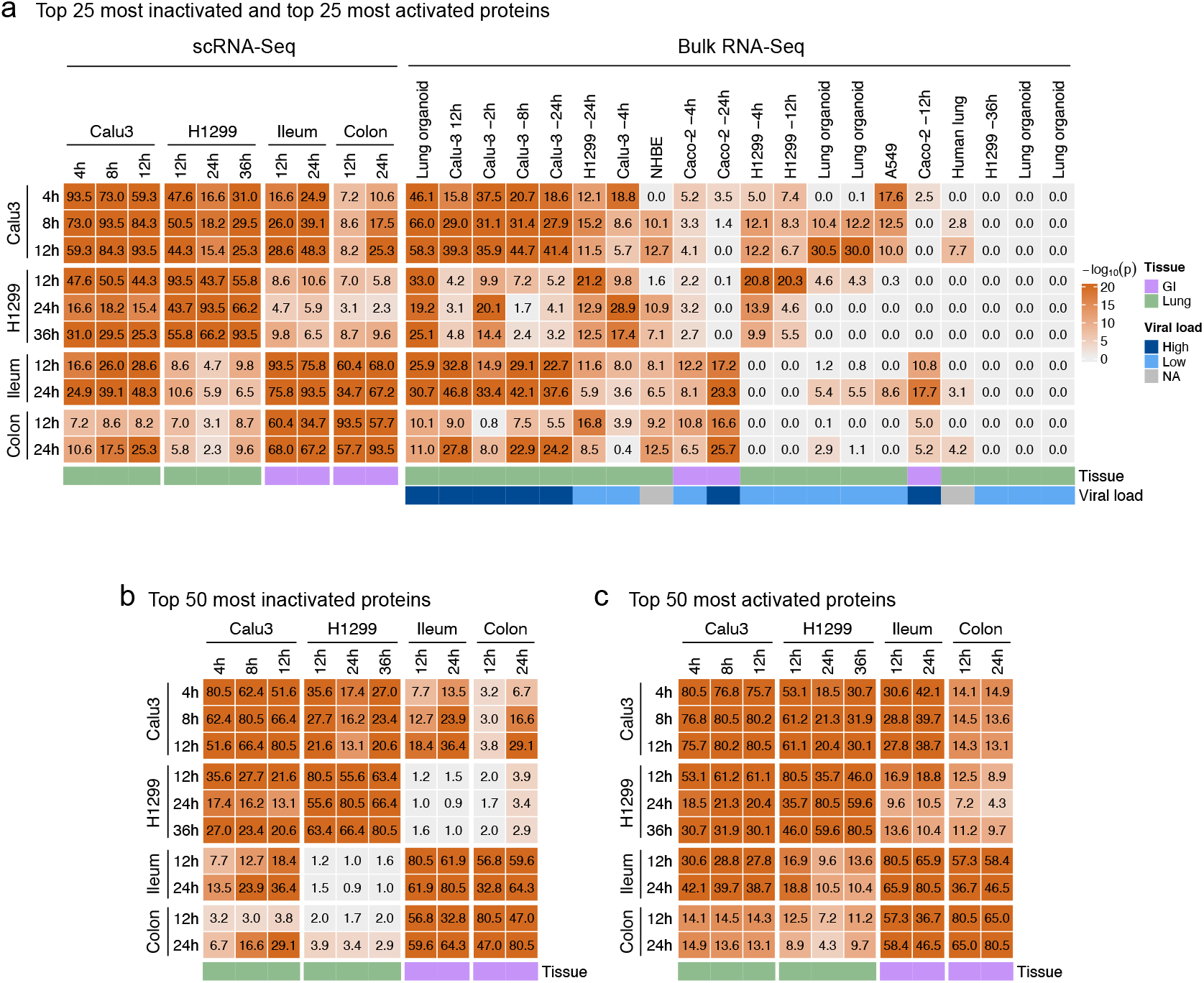
Conservation of VIPER-inferred Viral Checkpoint. **a**. Heatmap showing the conservation across single-cell and bulk-tissue samples. Results are expressed as -log_10_(p-value), estimated by the reciprocal enrichment of the 25 most activated and 25 most inactivated proteins in each signature using the aREA algorithm as implemented in the viperSimilarity function of the VIPER package. **b–c**. Conservation specifically for the top 50 most activated proteins (b) and most inactivated proteins (c) in response to SARS-CoV-2 infection between time points and models profiled at the single-cell level.

**Supplementary Figure 3.**
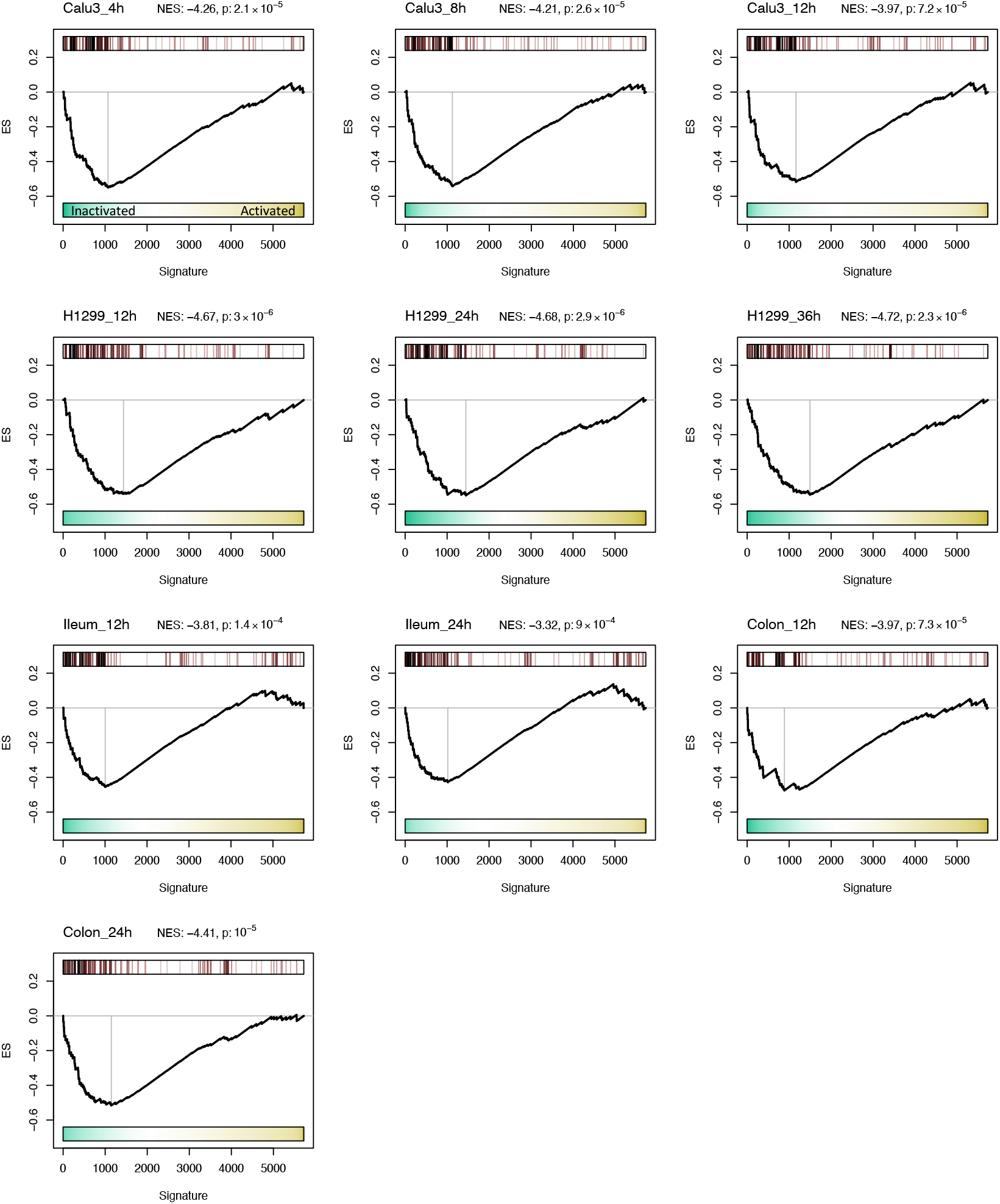
Enrichment of host factors known to physically interact with SARS-CoV-2 proteins on the host proteins differentially active in response to viral infection. GSEA showing the enrichment for the SARS-CoV-2 interacting proteins in the individual SARS-CoV-2 induced protein activity signatures. NES and p-values were estimated by one-tailed test and 1,000 permutations.

**Supplementary Figure 4.**
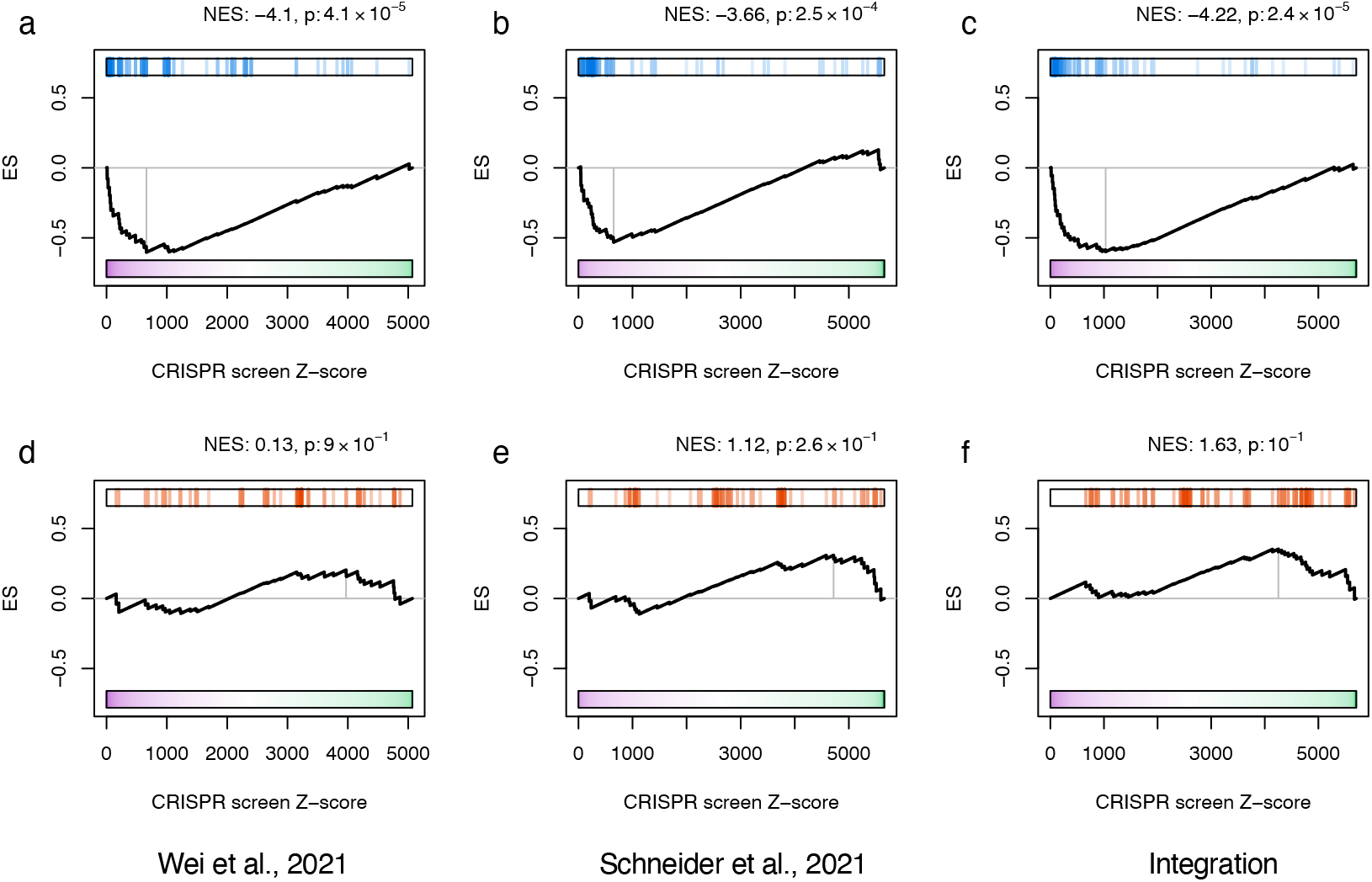
Enrichment of candidate SARS-CoV-2 infection MR proteins on host factors essential for SARS-CoV-2 infectivity. GSEA showing the enrichment of the top 50 most inactivated proteins in response to SARS-CoV-2 infection (inactivated candidate MR proteins) on the antiviral essential genes (a–c), but no enrichment of the top 50 most activated proteins in response to SARS-CoV-2 infection (activated candidate MR proteins) on the pro-viral essential genes (d–f), identified by 2 CRISPR screens (a, b, d and e) and their integration (c and f).

**Supplementary Figure 5.**
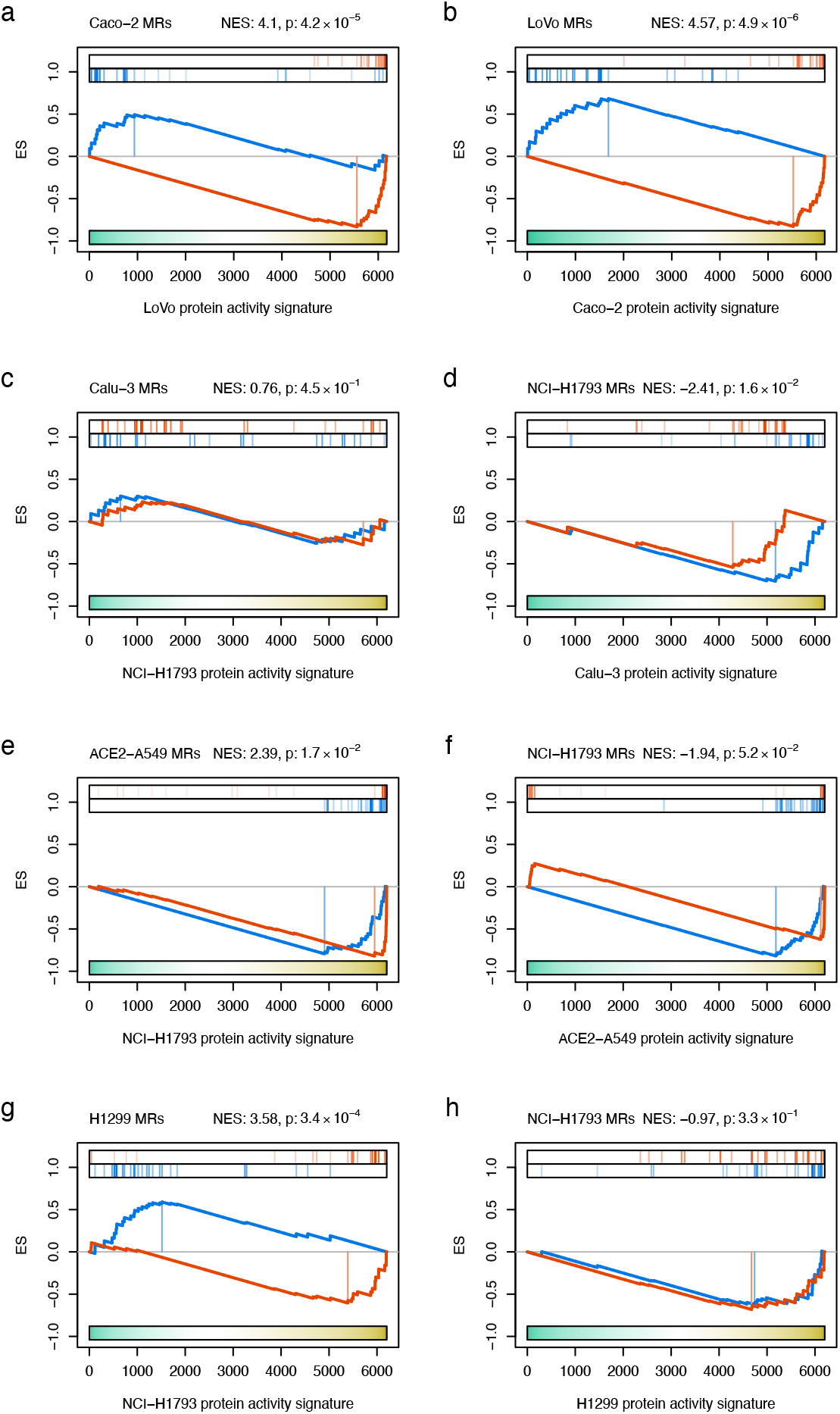
Conserved activity of MR proteins between cell line models susceptible to SARS-CoV-2 infection (Caco-2, Calu-3, ACE2-A549 and H1299) and the lineage context-matched cell lines included in the drug perturbation PANACEA resource (LoVo and NCI-H1793). **a**. GSEA for the enrichment of the Caco-2 top 25 most activated and top 25 most inactivated proteins in the LoVo protein activity signature. **b**. GSEA for the enrichment of the LoVo top 25 most activated and top 25 most inactivated proteins in the Caco-2 protein activity signature. **c**. GSEA for the enrichment of the Calu-3 top 25 most activated and top 25 most inactivated proteins in the NCI-H1793 protein activity signature. **d**. GSEA for the enrichment of the NCI-H1793 top 25 most activated and top 25 most inactivated proteins in the Calu-3 protein activity signature. **e**. GSEA for the enrichment of the ACE2-A549 top 25 most activated and top 25 most inactivated proteins in the NCI-H1793 protein activity signature. **f**. GSEA for the enrichment of the NCI-H1793 top 25 most activated and top 25 most inactivated proteins in the ACE2-A549 protein activity signature. **g**. GSEA for the enrichment of the H1299 top 25 most activated and top 25 most inactivated proteins in the NCI-H1793 protein activity signature. **h**. GSEA for the enrichment of the NCI-H1793 top 25 most activated and top 25 most inactivated proteins in the H1299 protein activity signature. Normalized Enrichment Score (NES) and p-value were estimated by two-tailed test and 1,000 permutations.

**Supplementary Figure 6.**
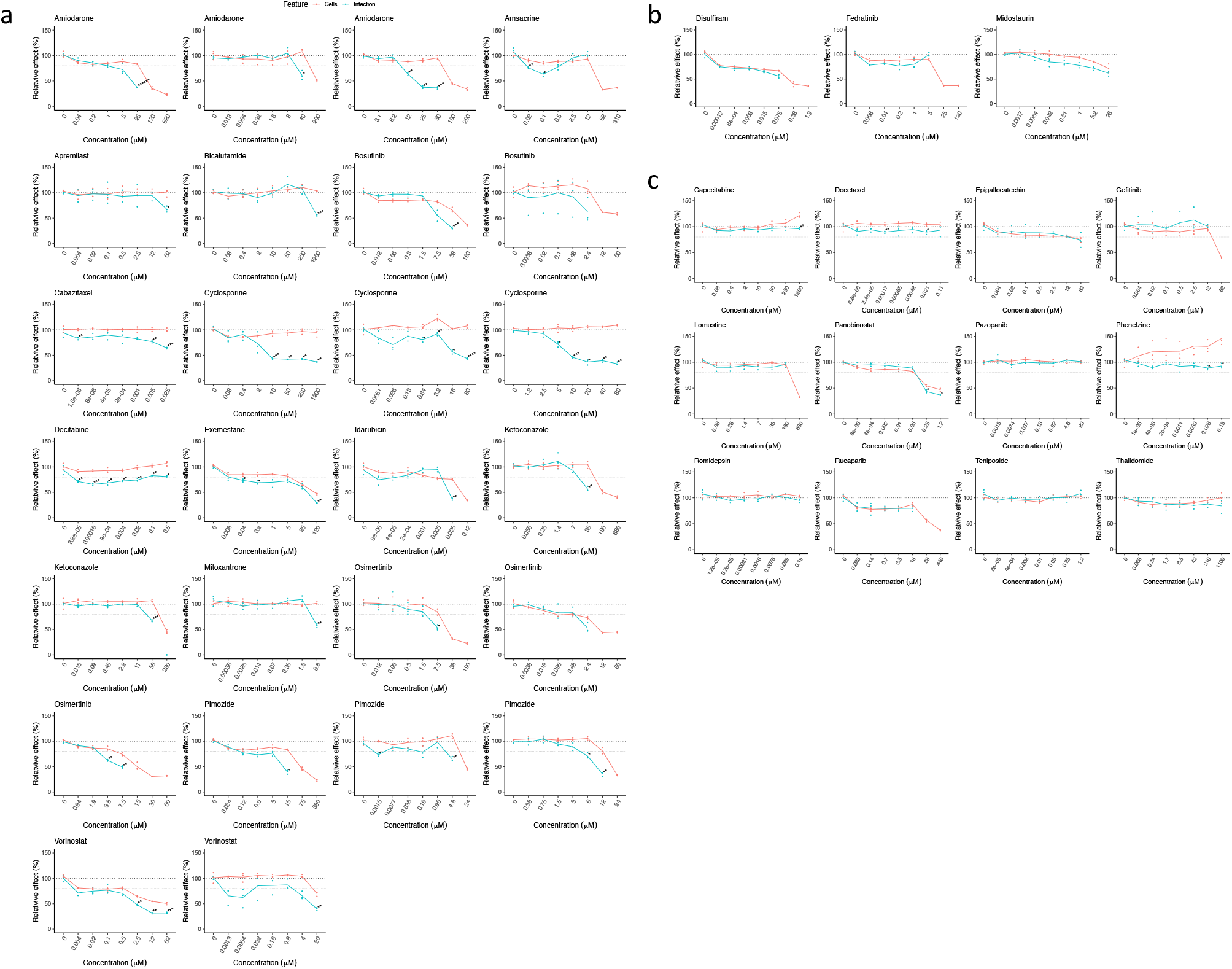
Experimental evaluation of the antiviral effect of FDA-approved drugs in Caco-2 cells. **a**. 15 of the 18 drugs predicted by ViroTreat showing significant antiviral effect (FDR *<* 0.05 and ≥ 20% infectivity decrease). **b**. 3 of the 18 drugs predicted by ViroTreat showing no significant antiviral effect. **c**. 12 drugs not significant by ViroTreat (p ≥ 0.01) selected as putative negative controls. The scatter-plots show the effect of each drug—SARS-CoV-2 infectivity shown in cyan and cell viability in red—relative to vehicle control (y-axis), assayed at different concentrations (x-axis) in triplicate. The lines indicate the average across replicates. * *p <* 0.05, ** *p <* 0.01, *** *p <* 0.001, **** *p <* 10^−4^, ****** *p <* 10^−6^, 1-tailed Student’s t-test, BC.

**Supplementary Figure 7.**
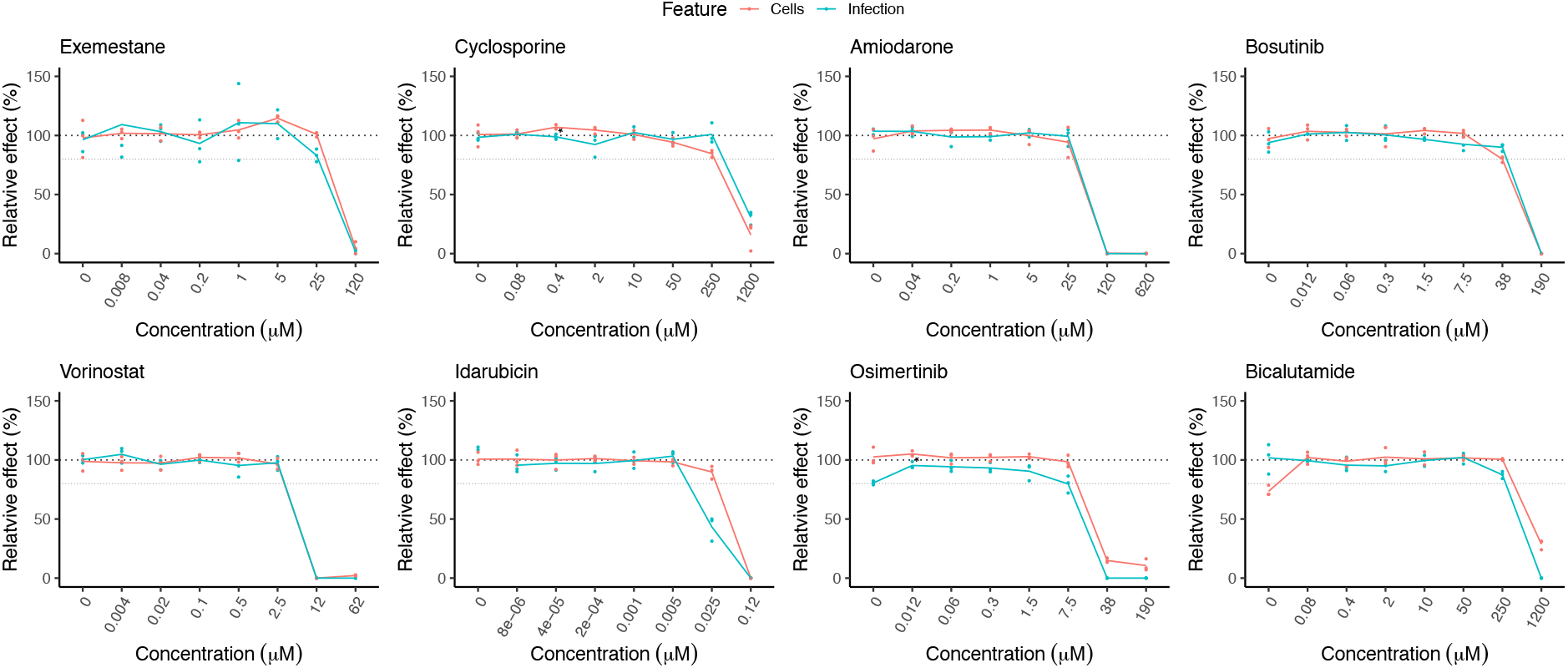
Experimental evaluation of 8 drugs, predicted by ViroTreat and showing the strongest SARS-CoV-2 antiviral effect in Caco-2 cells, for their effect on rotavirus infectivity. The scatter-plots show the effect of each drug—rotavirus infectivity shown in cyan and cell viability in red—relative to vehicle control (y-axis), assayed at different concentrations (x-axis) in triplicate. The lines indicate the average across replicates. * *p <* 0.05, 1-tailed Student’s t-test, BC.

**Supplementary Figure 8.**
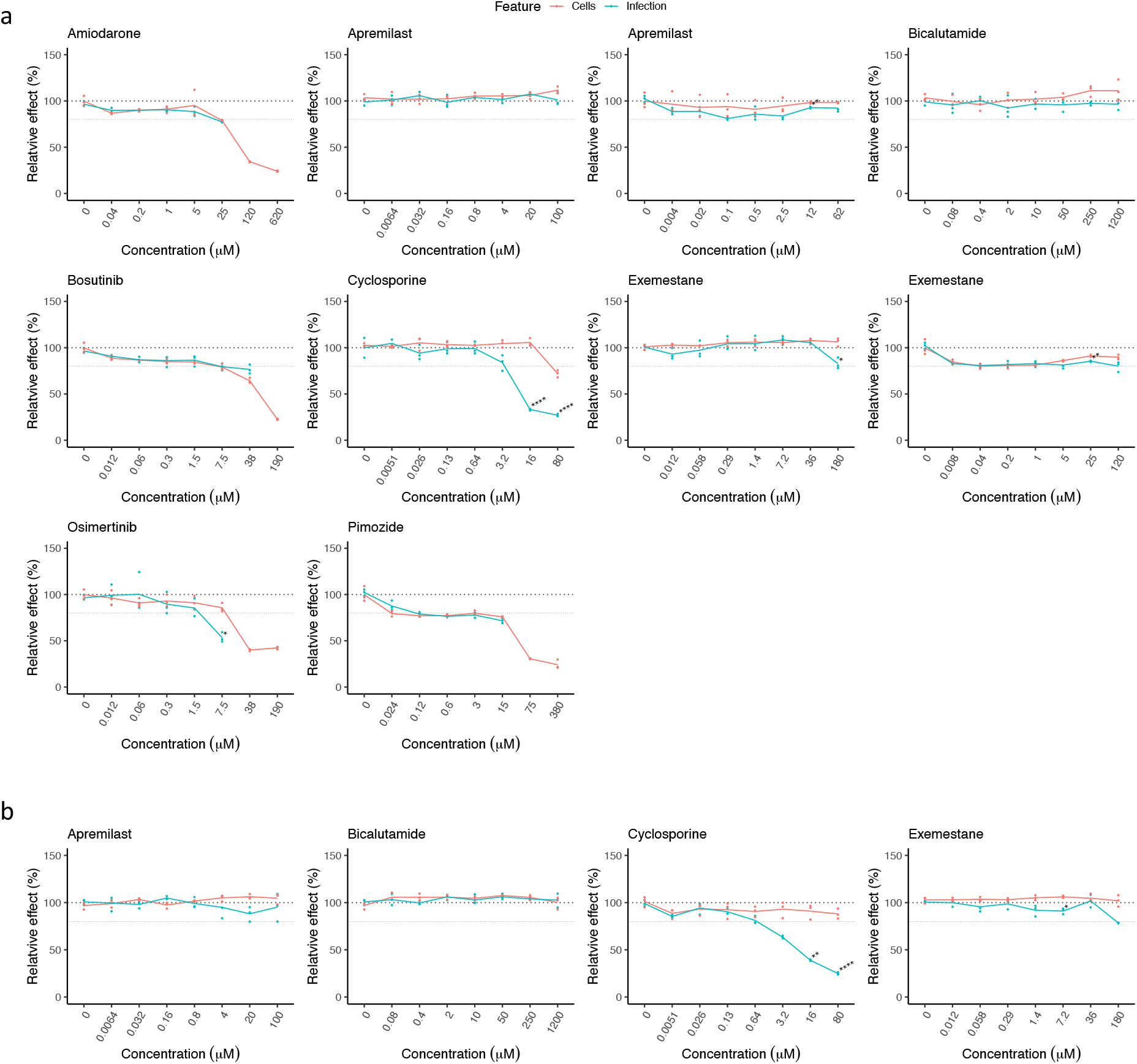
Experimental evaluation of the antiviral effect of FDA-approved drugs in lung adenocarcinoma cell lines. A set of drugs, predicted by ViroTreat for the GI context and with validated antiviral effect in Caco-2 cells were evaluated in Calu-3 (a) and A549-ACE2 (b) cells. The scatter-plots show the effect of each drug—SARS-CoV-2 infectivity shown in cyan and cell viability in red—relative to vehicle control (y-axis), assayed at different concentrations (x-axis) in triplicate. The lines indicate the average across replicates. * *p <* 0.05, ** *p <* 0.01, **** *p <* 10^−4^, 1-tailed Student’s t-test, BC.

**Supplementary Table 1.**
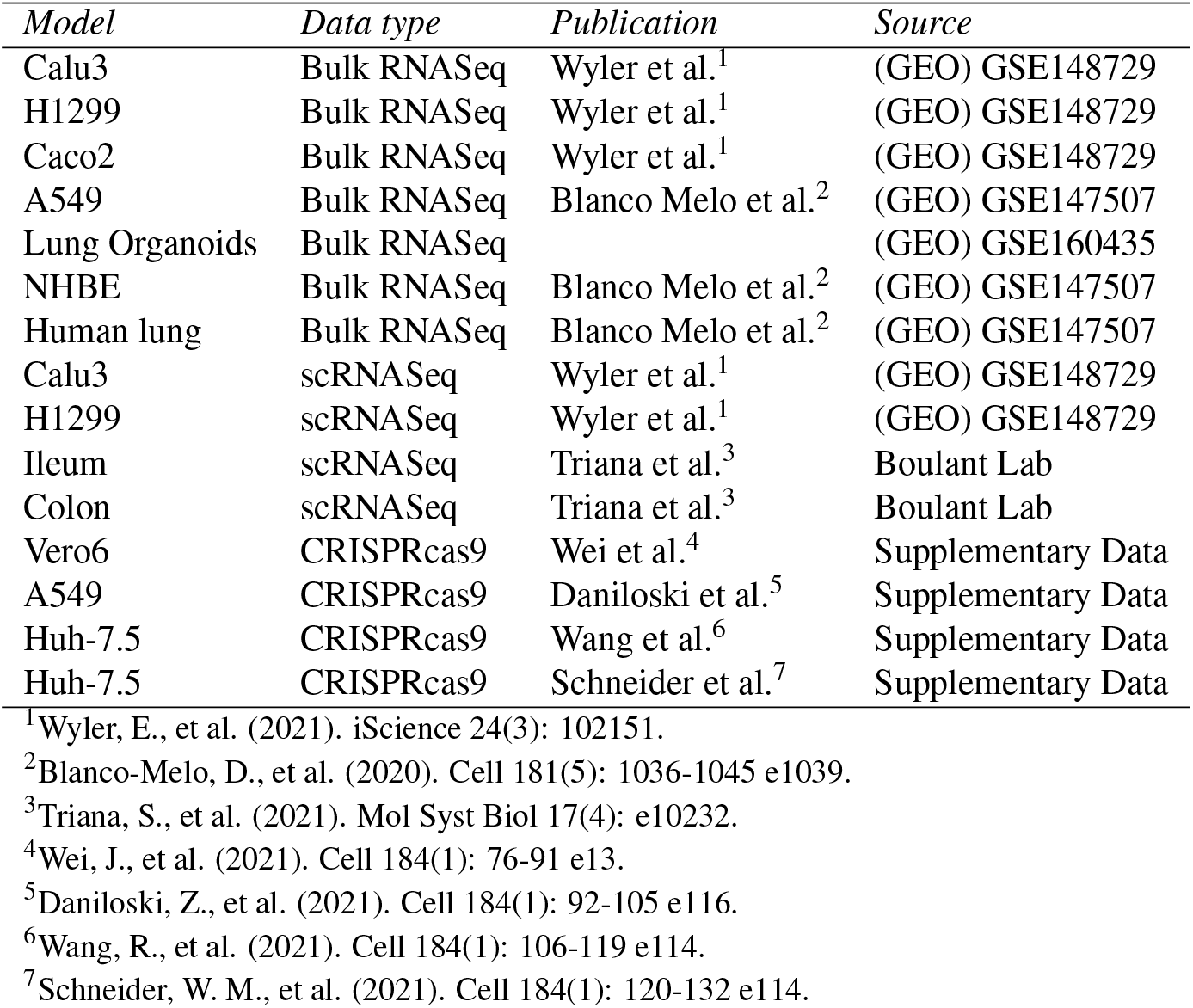
SARS-CoV-2 host cell RNA-Seq and scRNA-Seq datasets.

**Supplementary Table 2.** Drugs library, ViroTreat and focused validation screen results.

< See supplementary file Table-S2.xlsx >

**Supplementary Table 3.**
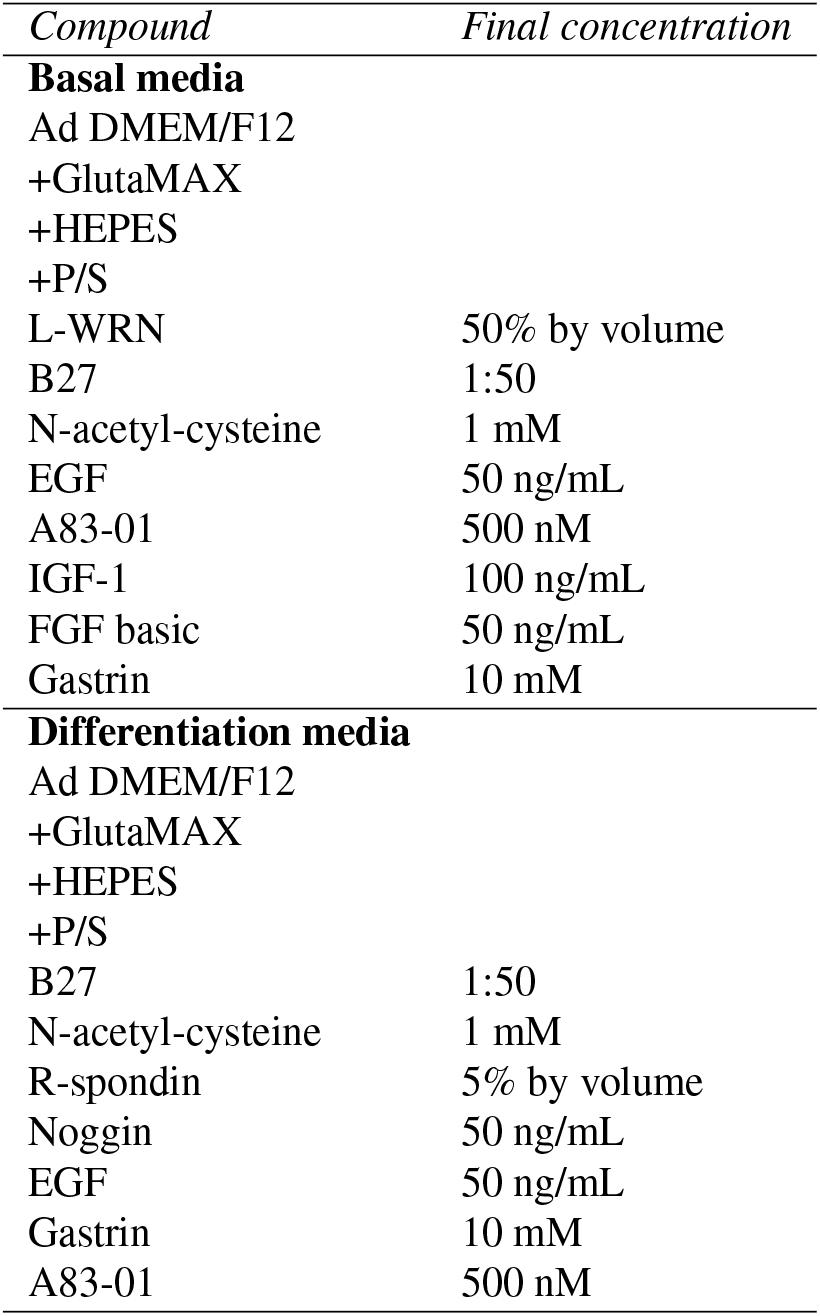
Organoids’ culture media.

| *Compound* | *Final concentration* |
| --- | --- |
| **Basal media** |  |
| Ad DMEM/F12 |  |
| +GlutaMAX |  |
| +HEPES |  |
| +P/S |  |
| L-WRN | 50% by volume |
| B27 | 1:50 |
| N-acetyl-cysteine | 1 mM |
| EGF | 50 ng/mL |
| A83-01 | 500 nM |
| IGF-1 | 100 ng/mL |
| FGF basic | 50 ng/mL |
| Gastrin | 10 mM |
| **Differentiation media** |  |
| Ad DMEM/F12 |  |
| +GlutaMAX |  |
| +HEPES |  |
| +P/S |  |
| B27 | 1:50 |
| N-acetyl-cysteine | 1 mM |
| R-spondin | 5% by volume |
| Noggin | 50 ng/mL |
| EGF | 50 ng/mL |
| Gastrin | 10 mM |
| A83-01 | 500 nM |

**Supplementary Table 4.** PCR primers.

| *Gene name* | *Species* | *Forward sequence* | *Reverse sequence* |
| --- | --- | --- | --- |
| HPRT1 | Human | cct ggc gtc gtg att agt gat | aga cgt tca gtc ctg tcc ata a |
| COV1 | SARS-CoV-2 | gcc tct tct gtt cct cat cac | aga cag cat cac cgc cat tg |

